# Large Language Models as Preprocessing Optimisers in Clinical Spectroscopy

**DOI:** 10.64898/2026.11.19.765453

**Authors:** Georgios Papadopoulos, Dimitar Georgiev, Álvaro Fernández-Galiana, Ho-Cheung Ng, Molly M. Stevens, Mauricio Barahona

## Abstract

Clinical spectroscopy has emerged as a powerful tool across biomedical research, with applications spanning from disease diagnosis and screening, to biomarker detection, therapeutic monitoring and intraoperative assessment. However, despite substantial progress, the analysis of spectroscopic data remains a major bottleneck, with current workflows still heavily reliant on trial and error, expert intuition and prior experience. Here, we introduce a multi-agent, large language model (LLM) framework for automated optimisation of preprocessing pipelines for spectral data. Our system combines a memory-derived information set, or *dossier*, with LLM-based agents that reason over previously evaluated preprocessing pipelines in order to propose, rank, and iteratively refine preprocessing strategies by following a multi-reward rule. Unlike exhaustive, brute-force search over a parameter space, the agent constructs candidates sequentially, enabling more targeted exploration. We benchmark our approach across 18 scenarios on six public clinical datasets against seven baselines, including random search, established optimisation methods and grid search. Our agentic optimisation achieved the highest classification accuracy in four out of six datasets. Ablation across LLM backbones and dossier configurations further shows that access to optimisation history is a key contributor to performance. These results demonstrate the potential of history-informed agents to enable more automated, adaptive and reproducible preprocessing pipelines in spectroscopy.

## 1 Introduction

Spectroscopic techniques, such as Raman spectroscopy (RS), Fourier transform infrared (FTIR) spectroscopy and diffuse optical/reflectance spectroscopy (DOS/DRS), enable rapid, non-destructive and label-free chemical analysis with minimal sample preparation (Butler et al., 2016; Baker et al., 2014; Bydlon et al., 2015). As such, they have emerged as powerful analytical tools for biomedical research and clinical medicine; with applications spanning from disease diagnosis and screening (Haka et al., 2005; Paraskevaidi et al., 2018; Giamougiannis et al., 2021b; Vališ et al., 2024), to biomarker detection (Qian et al., 2008; Plou et al., 2022), therapeutic monitoring (Cerussi et al., 2007; Cong et al., 2026) and intraoperative assessment (Jermyn et al., 2015; Hollon et al., 2020). Many of these applications involve (binary) classification of spectra into clinically relevant groups, such as disease states or tissue types. Despite substantial progress, however, robust analysis of spectroscopic data remains a major challenge, limiting the reproducibility, scalability and broader translation of spectroscopic methods into clinical practice (Baker et al., 2018; Coca-Lopez et al., 2025).

A critical component of spectroscopic data analysis is spectral preprocessing. Spectroscopic methods yield high-dimensional measurements that are often affected by various sources of non-specific variation, such as instrumental artefacts (e.g. cosmic-ray spikes), sample-derived backgrounds (e.g. fluorescence in RS, scattering-induced baselines in FTIR), experimental noise (e.g. detector and environmental noise), batch effects and sample preparation variability (Bocklitz et al., 2011; Isabelle et al., 2016; Butler et al., 2018; Vulchi et al., 2024). These can obscure biologically relevant spectral signatures and substantially degrade downstream analysis and predictive modelling. To mitigate these effects, spectroscopy workflows typically apply a series of preprocessing steps, such as despiking, smoothing, baseline correction and normalisation, to reduce non-specific variation and standardise collected measurements before analysis. While the impact of these steps is well documented (Bocklitz et al., 2011; Guo et al., 2021; Wahl et al., 2022), algorithm and hyperparameter selection remains largely reliant on manual trial and error, expert intuition and prior experience on comparable samples (Figure 1A).

**Figure 1:**
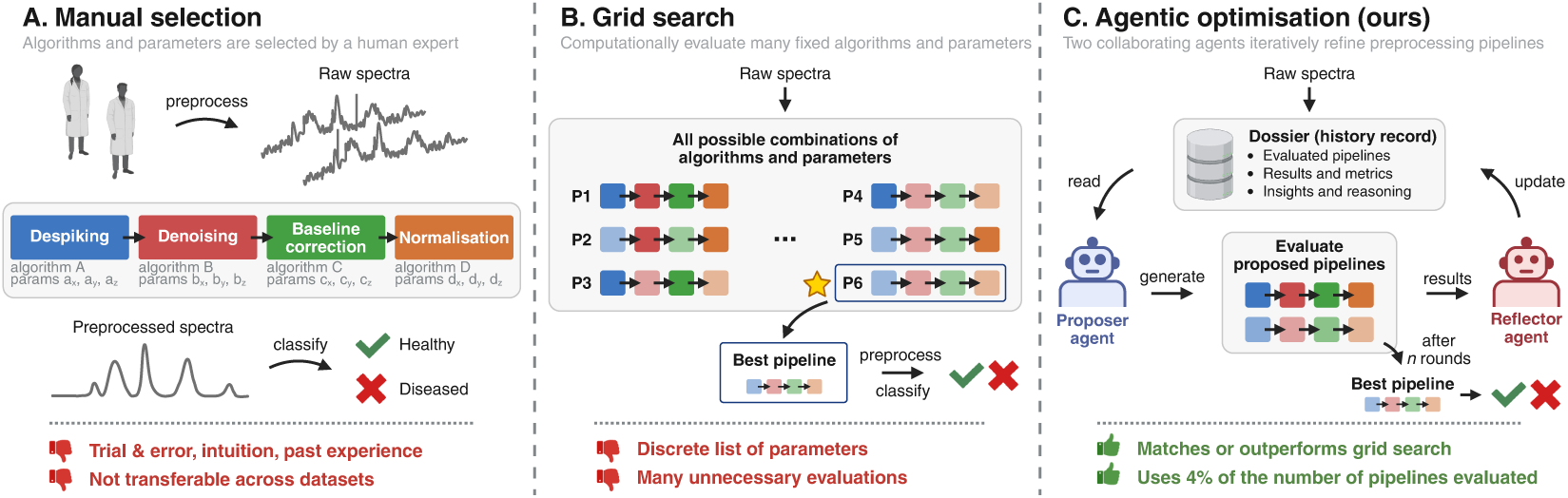
Comparative overview of preprocessing optimisation strategies in spectroscopy: manual selection (A), exhaustive grid search (B), and our proposed approach based on LLMs as optimisers (C).

A quantitative alternative is to search a predefined space of preprocessing pipelines and select configurations based on held-out performance. This is commonly achieved using exhaustive grid search over a selected set of algorithms and hyperparameters (Figure 1B) (Afseth et al., 2006; Lee et al., 2018; Storey & Helmy, 2019; Wahl et al., 2022; Kizilov et al., 2026; Han et al., 2026). However, exhaustive search is restricted to configurations specified in advance, can expend substantial computation on unpromising candidates, and may miss effective pipelines outside the search space. While more targeted strategies based on genetic algorithms (Jarvis & Goodacre, 2005; Bocklitz et al., 2011; Brunel et al., 2021), Bayesian optimisation (Babatunde et al., 2025) and statistical experimental design (Gerretzen et al., 2015) can reduce the number of candidates explored, identifying accurate and robust preprocessing pipelines across diverse datasets remains challenging.

Recently, LLM-based agents have emerged as optimisers that enable adaptive, feedback-informed search. Agents can propose candidates conditioned on past performance (Yang et al., 2024), synthesise directional feedback from the optimisation trace (Nie et al., 2024), route observations into the weights (Lv et al., 2025), and distil the accumulated history into a compact form (Ran et al., 2025). These capabilities enable optimisation strategies that iteratively interpret previous outcomes, revise the search trajectory, and explore complex spaces without exhaustively enumerating candidates. Consequently, agentic optimisation is increasingly being applied across scientific and engineering domains, from computational design to experimental and laboratory workflows (Boiko et al., 2023; Ma et al., 2024; Chen et al., 2025; Murphy, 2026).

Agentic approaches have also recently emerged in spectroscopy for spectral interpretation (Noh et al., 2026), literature parsing and analysis (Fu et al., 2025; Xie et al., 2026), instrument control (Lim et al., 2026), and experimental planning and automation (Rittig et al., 2026). However, their use for performance-driven optimisation of spectral preprocessing remains largely unexplored.

Here, we develop an LLM-based agentic system for automated preprocessing optimisation that iteratively proposes, evaluates and refines preprocessing pipelines by reasoning over the accumulated search history and adaptively exploring the space of algorithms and hyperparameters (Figure 1C).

### Contributions and findings

Our key contributions and findings are:

- We developed a multi-agent LLM-as-optimiser framework for automated optimisation of spectral preprocessing based on a *proposer* agent, a *reflector* agent, and a structured memory-derived information set over preprocessing pipelines, which we call *dossier*.
- We benchmarked two different LLM families and two parameter scales (8B and 70B parameters) against seven baseline methods across six clinical datasets (binary classification). Our agentic optimisation achieved the highest classification accuracy in four out of six datasets.
- We performed a range of ablation studies and empirically show that the history of the *dossier* is a key contributor to performance.

## 2 Problem formulation

### Preprocessing search space

In this work, we consider preprocessing pipelines *S* composed of four stages, namely:

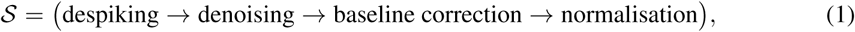

which operate on spectral measurements *x ∈* ℝ*^q^*, where *q* is the number of spectral channels, and are applied sequentially in the order specified in Equation 1, in line with established guidelines (Guo et al., 2021). These are preprocessing steps commonly used in spectral data analysis to remove cosmic-ray spikes, reduce noise, correct baseline variation, and normalise spectral intensity, respectively. Each stage *s ∈ S* comes with a corresponding library of methods *M_s_* to choose from, which also contains the identity ∅ (i.e. skip stage *s*). Each method *m ∈ M_s_* has a parameter domain Θ*_m_*, which is a product of integer ranges and real intervals subject to method-specific constraints. We write *θ ∈* Θ*_m_* for a particular setting of those parameters. For instance, the parameter domain Θ_SG_ of a Savitzky-Golay denoising filter can be defined as Θ_SG_ = {(*w, o, d*) *∈* N^3^ | *w* is odd}, where *w* specifies the window length, *o* specifies the polynomial order, and *d* specifies the derivative order (Savitzky & Golay, 1964).

The full action space *A* over *S* is thus given by the product:

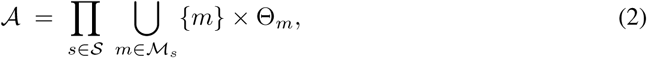

where each action *a* = [(*m_s_, θ_m_*)]*s∈S* defines a preprocessing pipeline that induces a transform *T_a_* : ℝ*^n×q^ →* ℝ*^n×q^′*, where *q^′^* is the number of spectral channels after preprocessing and *θ_m_∈*Θ. The action space *A* is typically infinite since Θ*_m_* contains real intervals. In our grid search experiments, we select a fixed set of discrete values for each real-valued Θ*_m_*.

### Pipeline evaluation

We focus on binary classification, a common framing in clinical spectroscopy and the one for which public cohorts are most widely available. We evaluate preprocessing strategies on clinical datasets *D* = (*X, y*) containing patient measurements *X ∈* ℝ*^n×q^* and binary labels *y ∈ {*0, 1*}^n^* (e.g. *healthy* vs. *diseased*). For a given pipeline *a*, we define a reward vector **r**(*a*) combining *ϕ* evaluation metrics *µ*_1_*,…, µ_ϕ_* as follows:

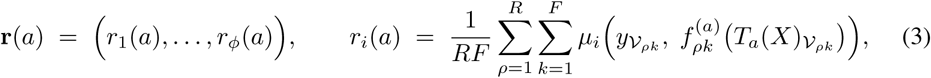

where, for repeat *ρ* and fold *k*, *I_ρk_* and *V_ρk_* are the disjoint index sets of the training and validation patients produced by stratified, patient-grouped *F*-fold cross-validation (CV), (*·*)*_I_ρk* denotes restric tion to those rows, and 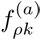 is the classifier fitted on (*T_a_*(*X*)*_I_, y_I_ρk*). Here, we set *ϕ* = 2 with *µ*_1_ := area under the receiver operating characteristic curve (AUC) and *µ*_2_ := sensitivity.

Three assumptions of Equation 3 carry through the paper. First, each repeat *ρ* draws a patient-level class-balanced subsample before train-test splitting, so every CV fold is balanced by construction and *µ_ϕ_* is not inflated by class imbalance. Second, every method in *Us M_s_* is a per-spectrum operator, so applying *T_a_* before splitting is equivalent to applying it within each fold and does not introduce information leakage. In contrast, feature scaling is not a per-spectrum operation and is therefore fitted separately within each fold. Third, fold assignments are seeded, making **r**(*a*) deterministic in *a*. Re-evaluating an action returns a bit-identical vector and yields no new information.

## 3 Proposed method

In our proposed setup, an LLM-based multi-agent system explores preprocessing algorithms and their parameter domains to iteratively design preprocessing pipelines and optimise classification performance. At each round, a *proposer* agent proposes candidate pipelines from a structured record of the evaluated history, denoted *dossier*, which includes previously top-performing pipelines, rejected pipelines, LLM predictions and data measures (Sections 3.1 and 3.2). Each proposal is checked against a registry of admissible parameter ranges before it enters the action space. Admitted candidates are evaluated by CV, and those whose aggregated gap to the top pipelines exceeds a fixed tolerance level are rejected based on a coherent risk operator racing rule over AUC and sensitivity (Section 3.3). A *reflector* agent, then collects performance and pipeline information from the round and summarises it into a short *invariant* carried into the next *dossier*. The *dossier* is then updated with the new evidence and used to guide subsequent proposals. This process continues until a fixed budget of evaluations is exhausted. The full process is summarised in Algorithm 1.

### Algorithm 1

Agentic-guided multi-reward racing

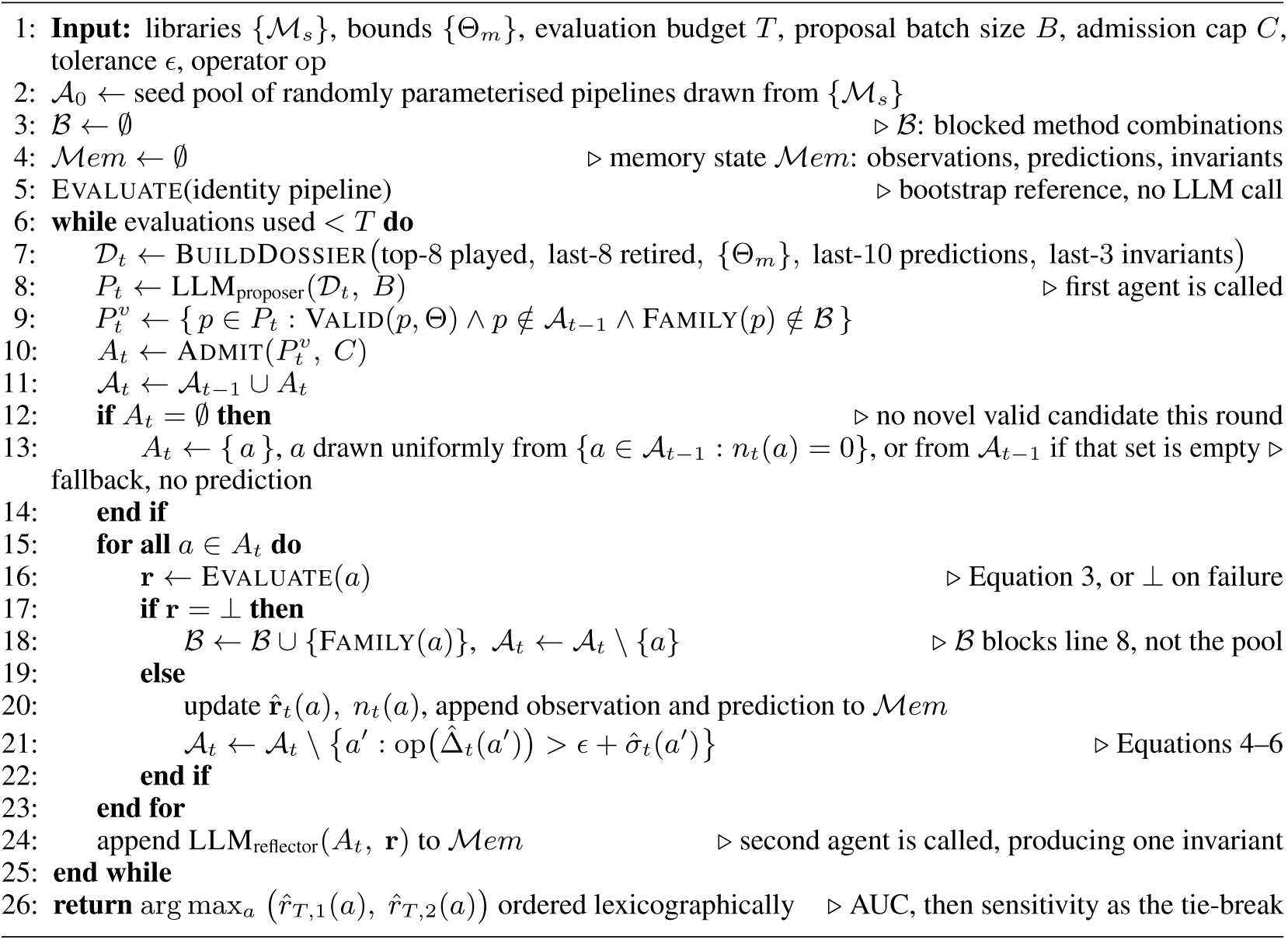

|  |  |  |
|---|---|---|
| 1: | **Input:** libraries $\{\mathcal{M}_s\}$, bounds $\{\Theta_m\}$, evaluation budget $T$, proposal batch size $B$, admission cap $C$, tolerance $\epsilon$, operator op |  |
| 2: | $\mathcal{A}_0 \leftarrow$ seed pool of randomly parameterised pipelines drawn from $\{\mathcal{M}_s\}$ |  |
| 3: | $\mathcal{B} \leftarrow \emptyset$ | ▷ $\mathcal{B}$: blocked method combinations |
| 4: | $\mathcal{M}em \leftarrow \emptyset$ | ▷ memory state $\mathcal{M}em$: observations, predictions, invariants |
| 5: | $\textsc{Evaluate}$(identity pipeline) | ▷ bootstrap reference, no LLM call |
| 6: | **while** evaluations used $< T$ **do** |  |
| 7: | $\mathcal{D}_t \leftarrow \textsc{BuildDossier}\big(\text{top-8 played},\ \text{last-8 retired},\ \{\Theta_m\},\ \text{last-10 predictions},\ \text{last-3 invariants}\big)$ |  |
| 8: | $P_t \leftarrow \text{LLM}_{\text{proposer}}(\mathcal{D}_t,\ B)$ | ▷ first agent is called |
| 9: | $P_t^v \leftarrow \{\, p \in P_t : \textsc{Valid}(p,\Theta) \wedge p \notin \mathcal{A}_{t-1} \wedge \textsc{Family}(p) \notin \mathcal{B} \,\}$ |  |
| 10: | $A_t \leftarrow \textsc{Admit}(P_t^v,\ C)$ |  |
| 11: | $\mathcal{A}_t \leftarrow \mathcal{A}_{t-1} \cup A_t$ |  |
| 12: | **if** $A_t = \emptyset$ **then** | ▷ no novel valid candidate this round |
| 13: | $A_t \leftarrow \{\, a \,\}$, $a$ drawn uniformly from $\{a \in \mathcal{A}_{t-1} : n_t(a) = 0\}$, or from $\mathcal{A}_{t-1}$ if that set is empty | ▷ fallback, no prediction |
| 14: | **end if** |  |
| 15: | **for all** $a \in A_t$ **do** |  |
| 16: | $\mathbf{r} \leftarrow \textsc{Evaluate}(a)$ | ▷ Equation 3, or ⊥ on failure |
| 17: | **if** $\mathbf{r} = \perp$ **then** |  |
| 18: | $\mathcal{B} \leftarrow \mathcal{B} \cup \{\textsc{Family}(a)\},\ \mathcal{A}_t \leftarrow \mathcal{A}_t \setminus \{a\}$ | ▷ $\mathcal{B}$ blocks line 8, not the pool |
| 19: | **else** |  |
| 20: | update $\hat{\mathbf{r}}_t(a),\ n_t(a)$, append observation and prediction to $\mathcal{M}em$ |  |
| 21: | $\mathcal{A}_t \leftarrow \mathcal{A}_t \setminus \big\{a' : \text{op}\big(\hat{\Delta}_t(a')\big) > \epsilon + \hat{\sigma}_t(a')\big\}$ | ▷ Equations 4–6 |
| 22: | **end if** |  |
| 23: | **end for** |  |
| 24: | append $\text{LLM}_{\text{reflector}}(A_t,\ \mathbf{r})$ to $\mathcal{M}em$ | ▷ second agent is called, producing one invariant |
| 25: | **end while** |  |
| 26: | **return** $\arg\max_a \big(\hat{r}_{T,1}(a),\ \hat{r}_{T,2}(a)\big)$ ordered lexicographically | ▷ AUC, then sensitivity as the tie-break |

### 3.1 Structure of the dossier

A key component of our approach is our *dossier*, which is inspired by the Optimization by PROmpting (OPRO)’s meta-prompt by Yang et al. (2024). We extend this idea by organising successful and unsuccessful pipelines, performance comparisons, diagnostic feedback and task constraints into a structured record. The aim is to help agents identify promising configurations, learn from previous evaluations and avoid repeating unsuccessful choices. The *dossier* is built from scratch each round and captures preprocessing pipelines have been proposed, past predictions, reasoning, and evaluation history. These are organised into eight sections as follows: the task and remaining budget (§1); the top-ranked evaluated pipelines with their scores (§2); the pipelines that have been rejected/retired together with the reason for rejection/retirement (§3, see following sections); the top-performing methods *m_s_* for each stage *s*, together with the *reflector*’s *invariants* (§4, see Section 3.2); the parameter domains Θ*_m_* (§5); model predictions paired with realised values (§6), spectral diagnostics for the current best (§7, computed on the training set); and the output contract (§8), which declares the method libraries *M_s_*, the JSON schema each proposal must satisfy, and the rules constraining it. Figure 5 and Table 2 in Appendix A provide further information about the structure of the *dossier* and examples for each section. §7 is the only section containing data-derived measurements rather than search bookkeeping, covering class separability, shape preservation and peak structure (see Table 3 in Appendix A). Three *dossier* sections are capped to keep the *dossier* bounded in size: §2 contains the eight best-scoring pipelines, §3 contains the eight most recent retirements, and §6 contains the ten most recent predictions.

### 3.2 The proposer and reflector agents

At each round *t*, a dossier *D_t_* is built and passed to the *proposer* LLM agent, which generates a batch of *B* pipeline candidates, *P_t_*, constructed using RamanSPy (Georgiev et al., 2024). Unlike in the original OPRO implementation by Yang et al. (2024), the generation is not trusted and each generated candidate pipeline *p* is parsed and checked against the parameter domains Θ. If a violation is detected, the candidate *p* is rejected with a logged reason, so that an invalid configuration can never enter the active set *A_t_ ⊆ A* as a valid pipeline *a*. Up to *C* pipelines are admitted to *A_t_* each round, which are individually evaluated using patient-level *F*-fold cross-validation, based on the reward defined in Equation 3. Note that the *proposer* is used before the pipelines are scored, so it never sees the outcome of its own batch directly.

After each pipeline in the active set is evaluated, our second, *reflector* LLM agent receives a one-line summary for each admitted pipeline, containing the predicted and observed AUC and sensitivity, the signed error, and a justification the *proposer* had offered. The use of a *reflector* agent in our agentic system is inspired by the work of Nie et al. (2024), who showed that synthesising directional feedback can improve the robustness and efficiency of agentic optimisation. In our framework, the *reflector* summarises the outcomes of each optimisation round and provides feedback to the *proposer* to guide subsequent pipeline proposals. At the end of each round, the *reflector* model is asked to produce a single statement of at most twenty words, which we call an *invariant*, capturing what the *proposer* should not change on the next round. To limit error propagation through free-text memory, we constrain the generation as follows: outputs are capped at 256 tokens, reduced to the first non-empty line, and discarded if they begin with deliberative markers (e.g. *okay*, *let’s*), or return the sentinel `NONE` for inconclusive batches.

A persistent memory store *Mem* retains the reward vector measured for every evaluated pipeline, every predicted score paired with the actual score, and every accepted *invariant*. It reaches the model through exactly two reads, both ordered by recency rather than relevance: the ten most recent prediction and outcome pairs populate §6, and the three most recent *invariants* populate §4. The model never queries the store *Mem*, so an *invariant* is the only channel by which a conclusion from round *t* reaches round *t* + 1 as language rather than as a number (see Figures 5 and 6 in Appendix A). At the end of each round, we clear each *dossier* section and re-run the search, so the contribution of the accumulated evidence is measured rather than assumed. The search continues until the predefined budget *T* of CV evaluations is reached. The number of rounds is an outcome, since a round contributes between zero and *C* evaluations.

### 3.3 Multi-reward racing

Each round, pipelines whose gap to the best-performing candidates is large enough are retired. We quantify this using the per-reward gap to the incumbent leaders, which we define as follows, inspired by Dann et al. (2023):

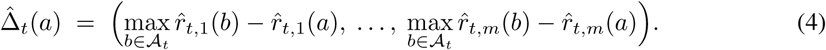

We aggregate the calculated per-reward gaps Δ^*t*(*a*) using an operator op : ℝ*^m^ →* ℝ. This must be a coherent risk measure, meaning monotone, positively homogeneous, sub-additive and translation invariant, and we use the conditional value-at-risk family,

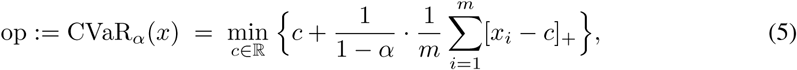

which recovers the mean at *α* = 0 and the maximum as *α →* 1. Here *c* is an auxiliary threshold of the Rockafellar-Uryasev variational form (Rockafellar & Uryasev, 2000), eliminated by the minimisation. The objective is convex and piecewise linear in *c*, with breakpoints at the entries of *x*, so minimising over *c* is exact. Choosing op = max retires an action as soon as any single reward shows it to be suboptimal, while op = mean requires the average gap to be large.

In our setup, an action is retired when its aggregated gap exceeds a fixed tolerance *ɛ* plus its own empirical dispersion *σ*^*_t_*(*a*), i.e.:

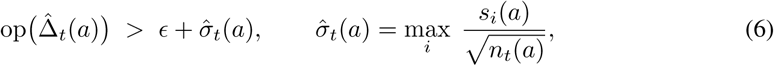

where *s_i_*(*a*) := *sd*(*r_i_*(*a*)) denotes the sample standard deviation of the observed rewards *r_i_*(*a*), computed over the *n_t_*(*a*) evaluations of pipeline *a* up to round *t*. An action with *n_t_*(*a*) *<* 2 has no estimable spread and is given a prior *σ*_0_ = 0.05, set on the scale of the spread between pipelines rather than that of the reward range. Since evaluation is deterministic across all our experiments, no action is evaluated more than once, so *n_t_*(*a*) *∈* {0, 1} and thus 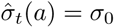 and 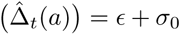 for all *a* and *t*.

### 3.4 Failed evaluations are missing data

A pipeline may crash, time out, or produce a non-finite metric (e.g. when using an unstable baseline fit). Such an outcome provides limited information about the quality of the pipeline and is thus recorded as a blank record *⊥* rather than as a score, to avoid assigning a non-running pipeline a higher rank (e.g. 0.5) than a pipeline that ran but scored below chance. On *⊥*, we record no reward, admit no memory observation, and drop the action. Its method combination enters the list of blocked actions *B*, and no further parameterisation of it may be proposed, since a combination that fails numerically usually fails across its parameter range. The block applies to admission and not to the pool.

## 4 Experimental setup

### Datasets

We performed experiments on six publicly available clinical datasets, spanning three spectroscopic techniques, two disease areas and three cohort sizes: (i) surface-enhanced Raman spectroscopy (SERS) measurements from plasma samples from healthy and ovarian cancer patients (Paraskevaidi et al., 2018), (ii-v) attenuated total reflection (ATR)-FTIR measurements from plasma (ii, iv) and serum (iii, v) samples from healthy and ovarian cancer patients (treated and non-treated with chemotherapy) (Giamougiannis et al., 2021a;b), and (vi) DRS measurements from skin for dengue screening (Poh et al., 2020). Further information about cohort sizes and spectral acquisition is provided in Table 4 in Appendix B. Each dataset was re-derived from the deposited raw files, prior to any preprocessing. When multiple measurements were available for a patient, these were averaged before preprocessing and classification, so measurements from the same patient cannot appear in both the training and test sets.

### Evaluation protocol

We performed experiments on six dataset combinations of modality, specimen and task, using three classifier families – a radial basis function (RBF)-kernel, support vector machine (SVM), partial least squares discriminant analysis (PLS-DA) and *ℓ*_2_-regularised logistic regression (LogReg). To assess generalisation, we benchmarked all optimisation methods on independent held-out test sets using patient-grouped 80/20 train–test splits (R=2 repeats). Optimisation and pipeline selection, including the type of classifier to use for final comparisons, were restricted to the training data, and final performance was assessed using held-out AUC. Ablation studies and optimisation diagnostics used R=5 repeats of 5-fold CV to characterise optimisation behaviour and component contributions. To allow meaningful comparisons, all methods were evaluated using the same metrics and CV splits.

### Agentic optimisation

For each of the 18 scenarios, we ran three independent agentic campaigns and performed preprocessing optimisation using three different LLM backbones for the *proposal* and *reflector* models: `deepseek-r1:8b`, `llama3.1:8b` and `llama3.1:70b`. All LLMs were run locally using a temperature of 0 over 2,160 evaluations across the 18 scenarios. Each campaign was run with a budget of 30 iterations, proposal batch size *B* = 10, admission cap *C* = 5, op = mean, *ɛ* = 0.01, *σ*_0_ = 0.05.

### Baselines

We benchmark our agentic optimisation against no preprocessing (**raw**), a full exhaustive evaluation of all pipelines over a predefined discrete parameter grid (**grid brute force**), and five baseline optimisers: random search, which samples pipeline configurations at random (**random**); sequential stage-wise optimisation, where each preprocessing stage is optimised in turn and the best choice is fixed before proceeding to the next (**greedy stepwise**); the Tree-structured Parzen Estimator (**TPE**) and genetic algorithms (**GA**), previously investigated for spectral preprocessing by Brunel et al. (2021); and Gaussian process-based Bayesian optimisation (**GP-BO**), as applied by Babatunde et al. (2025). Baseline optimisers were built using Optuna (Akiba et al., 2019), using the following settings: **random** draws each stage’s method uniformly (identity included) and its parameters uniformly within their ranges; **TPE** uses a tree-structured conditional encoding with multivariate, grouped sampling and 10 random start-up trials; **GP-BO** uses a Matérn-5/2 kernel with log expected improvement, a noise-free objective and 10 random start-up trials; and **GA** is a single-objective NSGA-II with population 20 as in Brunel et al. (2021), binary-tournament selection, uniform crossover (*p* = 0.9) and random-reset mutation. Each optimiser was run with 10 random seeds per scenario during inner cross-validation, and results were averaged across seeds.

### Search space

The agentic methods and budget-matched optimisers (random search, TPE, GA, and GP-BO) use the same preprocessing methods and parameter bounds, with a budget of 120 pipeline evaluations per run. These methods explore continuous parameter ranges, whereas exhaustive grid search evaluates predefined discrete values. To assess how reliably agents follow the specified stage definitions, their proposals are checked for valid methods and parameter bounds, but stage membership is not enforced (e.g., ASLS may be proposed as a denoiser). Post-hoc analysis showed that such cross-stage placements occurred in only 1.4–1.6% of admitted proposals from the two 8B models, none from `llama3.1:70b`, and just 1 of the 108 best selected pipelines by individual models across all 18 scenarios and two repeats (Table 17 in Appendix D). Note that the number of pipelines considered by *grid brute force* grows multiplicatively with the number of preprocessing algorithms and parameter choices: we perform up to *∼* 2,500 evaluations for each scenario, compared with only *∼* 45 for *greedy stepwise*.

## 5 Results

### 5.1 Bad preprocessing is worse than no preprocessing

First, we examine the effect of preprocessing on downstream classification. Figure 2 shows the distribution of classification performance across all preprocessing pipelines evaluated in the exhaustive *grid brute force* search and all 18 scenarios. We make the following two observations. First, the raw-data baseline is not a lower bound on performance: depending on the scenario, between 0.7% and 65% of the tested pipelines perform worse than the identity pipeline (i.e. no preprocessing). Hence, preprocessing is not guaranteed to improve classification performance and can just as readily degrade it. Second, dispersion generally decreases as mean performance increases, with the noisiest evaluations concentrated in the lower-performing pipelines, which an adaptive search should discard.

**Figure 2:**
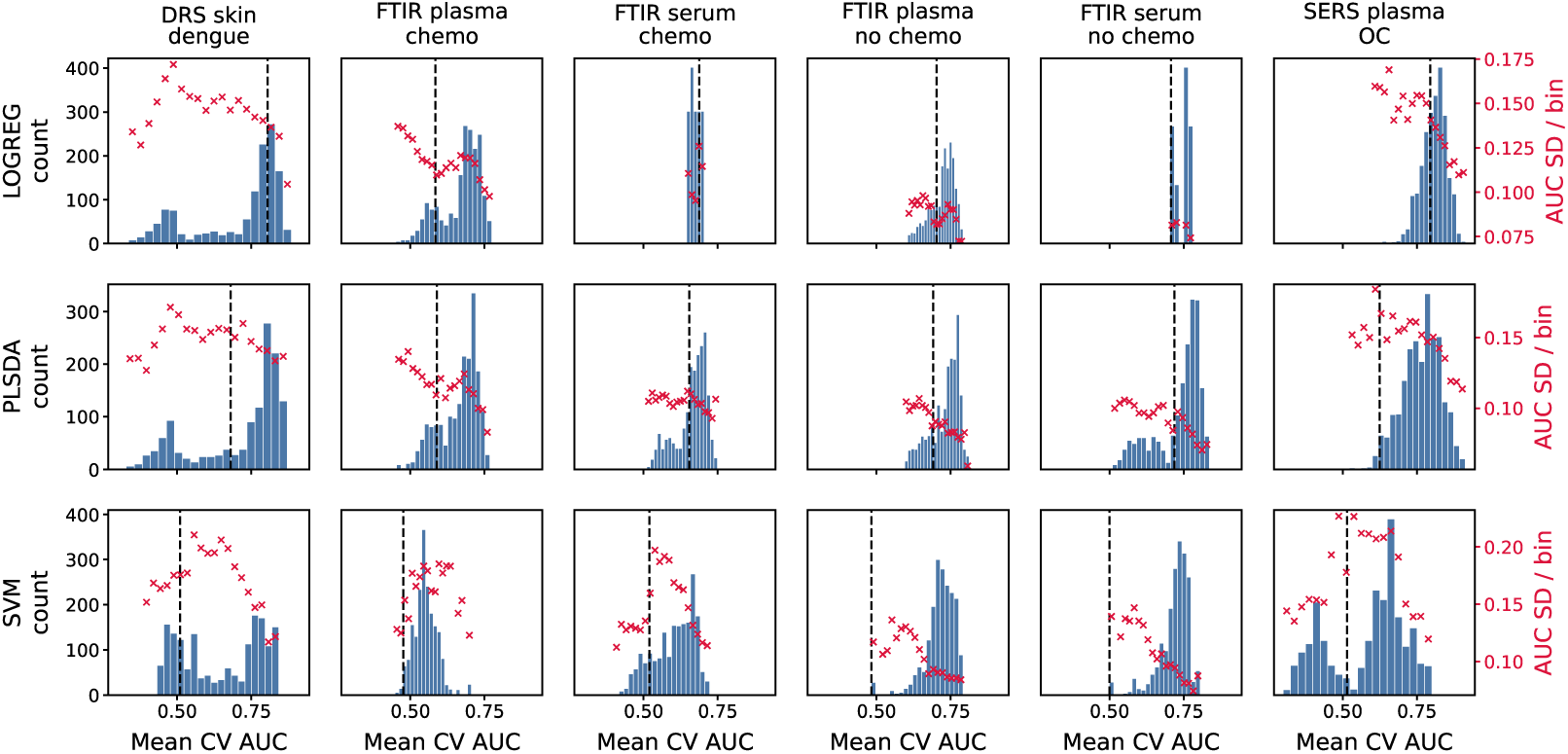
Distribution of classification performance (in-sample) across *∼* 2, 500 preprocessing pipelines evaluated during *grid brute force* search for each of the 18 scenarios considered. Dashed lines indicate performance on raw data. Red crosses show the standard deviation of AUC values for each bin.

### 5.2 Agentic optimisation outperforms grid search and established optimisers in four out of six clinical datasets

Building on these results, we benchmark our agentic system against the three baseline methods on all six datasets. Table 1 summarises the results, showing the out-of-sample classification performance of the best preprocessing pipeline selected by each optimisation strategy for each dataset (extended results are provided in Figures 7 and 8 and Tables 5–8 in Appendix C).

**Table 1:**
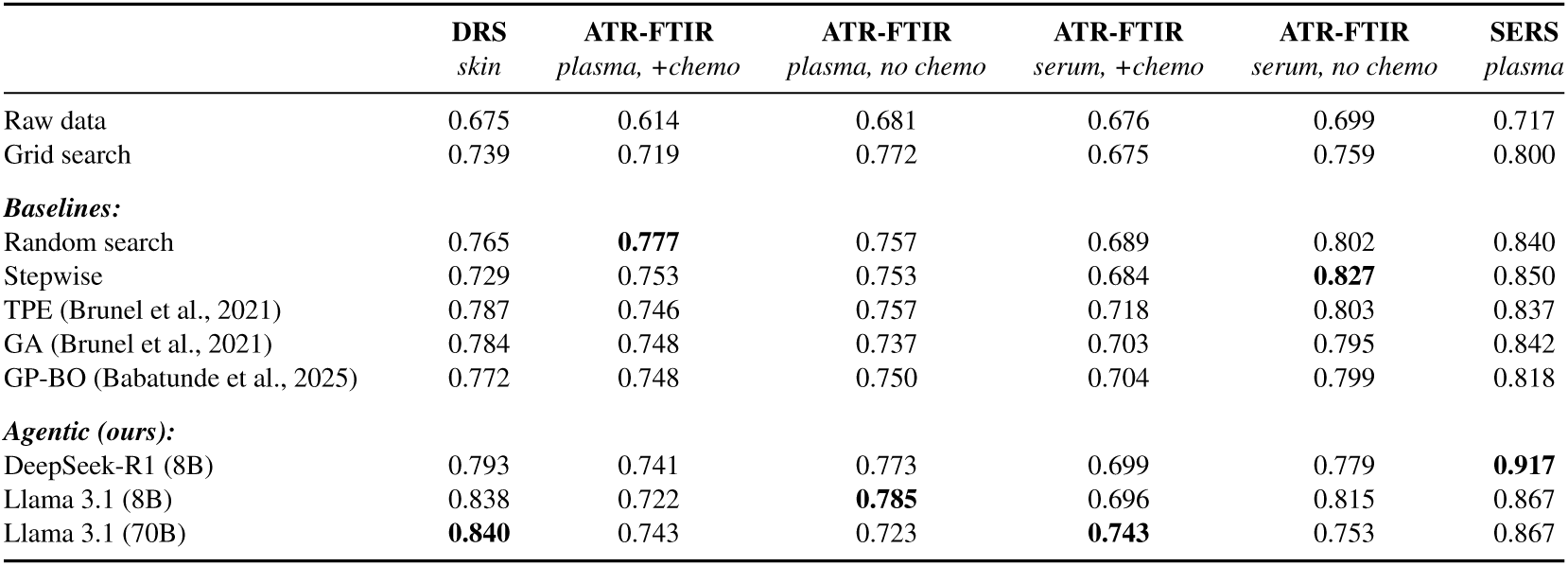
Benchmark results. Out-of-sample classification performance across the six datasets. Reported values show the held-out AUCs of the best pipelines and classifiers selected through internal cross-validation, averaged over two repeats. Best results are shown in bold.

|  | **DRS**<br>*skin* | **ATR-FTIR**<br>*plasma, +chemo* | **ATR-FTIR**<br>*plasma, no chemo* | **ATR-FTIR**<br>*serum, +chemo* | **ATR-FTIR**<br>*serum, no chemo* | **SERS**<br>*plasma* |
|---|---|---|---|---|---|---|
| Raw data | 0.675 | 0.614 | 0.681 | 0.676 | 0.699 | 0.717 |
| Grid search | 0.739 | 0.719 | 0.772 | 0.675 | 0.759 | 0.800 |
| ***Baselines:*** |  |  |  |  |  |  |
| Random search | 0.765 | **0.777** | 0.757 | 0.689 | 0.802 | 0.840 |
| Stepwise | 0.729 | 0.753 | 0.753 | 0.684 | **0.827** | 0.850 |
| TPE (Brunel et al., 2021) | 0.787 | 0.746 | 0.757 | 0.718 | 0.803 | 0.837 |
| GA (Brunel et al., 2021) | 0.784 | 0.748 | 0.737 | 0.703 | 0.795 | 0.842 |
| GP-BO (Babatunde et al., 2025) | 0.772 | 0.748 | 0.750 | 0.704 | 0.799 | 0.818 |
| ***Agentic (ours):*** |  |  |  |  |  |  |
| DeepSeek-R1 (8B) | 0.793 | 0.741 | 0.773 | 0.699 | 0.779 | **0.917** |
| Llama 3.1 (8B) | 0.838 | 0.722 | **0.785** | 0.696 | 0.815 | 0.867 |
| Llama 3.1 (70B) | **0.840** | 0.743 | 0.723 | **0.743** | 0.753 | 0.867 |

Across the six datasets, our agentic approach achieved the highest mean held-out AUC in four settings and outperformed the best of the three established optimisers (TPE, GA and GP-BO) in five. Compared with the best conventional baseline overall, the largest gains were observed for SERS plasma (0.917 vs. 0.850) and DRS skin (0.840 vs. 0.787), followed by ATR-FTIR serum with chemotherapy (0.743 vs. 0.718) and plasma without chemotherapy (0.785 vs. 0.772). Stepwise optimisation led on serum without chemotherapy (0.827 vs. 0.815), although our agents still outperformed the three established optimisers (best: 0.803). Random search led on plasma with chemotherapy (0.777 vs. 0.743), where the best established optimiser also performed slightly better (0.748).

Averaged across the 18 scenarios, the three agentic campaigns achieved held-out AUCs of 0.745 (`deepseek-r1:8b`), 0.750 (`llama3.1:8b`) and 0.753 (`llama3.1:70b`), compared with 0.631 for raw data, 0.726 for the *grid brute force*, 0.725 for *greedy stepwise*, 0.733 for *random method*, 0.739 for *TPE*, 0.733 for *GA*, and 0.734 for *GP-BO*. The three agentic campaigns outper-formed the mean of the three benchmark optimisation methods (TPE, GA and GP-BO) with mean out-of-sample AUC improvements of +0.010 *±* 0.010, +0.015 *±* 0.011 and +0.018 *±* 0.010 per scenario, respectively. All three agentic campaigns also outperformed *random* search, in 14 of the 18 scenarios, with mean out-of-sample AUC improvements of +0.012 *±* 0.014, +0.018 *±* 0.014 and +0.020 *±* 0.012 per scenario, respectively.

The agentic search required substantially fewer evaluations than *grid brute force*, with roughly 120 evaluations per campaign and scenario, compared with *∼* 2,500 pipelines for *grid brute force*. This resulted in runtimes of approximately 25–40 minutes per scenario on average for our three agentic optimisers compared to about 4 hours for *grid brute force* on average per scenario tested (see Table 9 in Appendix C). In comparison, conventional optimisers (i.e. TPE, GA and GP-BO) took about 6–7 minutes, but did not achieve the highest classification accuracy in any of the six datasets. We believe this trade-off may be worthwhile in many clinical applications, where even modest improvements in classification performance can justify additional optimisation time.

### 5.3 Preprocessing proposals improve as optimisation history accumulates

Next, we examine how proposal quality changes as optimisation history accumulates. Figure 3 shows the optimisation traces of our three agentic campaigns for a representative scenario [SERS (plasma, LogReg)], depicting the best proposals at each round (dotted lines) and the running best (solid lines). In this example, the agentic search with `deepseek-r1:8b` discovers the best preprocessing pipeline, reaching an AUC of 0.929 at round 16 and outperforming *greedy stepwise* optimisation (0.893). However, note that its performance varies considerably across rounds, with many proposals falling well below the best pipeline found so far. In contrast, the two `llama3.1` campaigns improve more steadily but plateau at AUCs of 0.889 (8B) and 0.880 (70B). The best preprocessing pipelines derived by each of the three agentic campaigns on this scenario are given in Table 10 in Appendix C.

**Figure 3:**
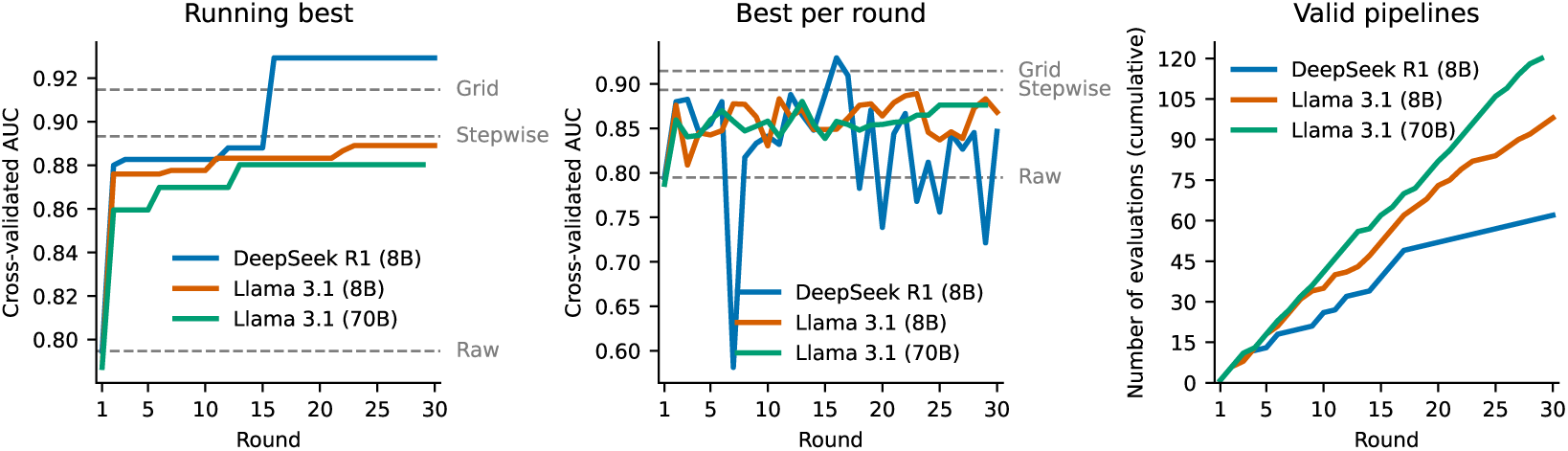
Classification performance (in-sample) across 30 rounds of preprocessing optimisation guided by agentic search on a selected scenario [SERS (plasma, LogReg)].

Extended analysis across all 18 scenarios shows that proposal quality generally improves as optimisation progresses, with a mean AUC of LLM-proposed pipelines increasing from 0.686 in the first third of the campaigns to 0.720 in the final third. In fact, performance improves in 16 out of the 18 scenarios (*p* = 0.001, sign test), with a positive correlation with round index in 14 of 18 (*p* = 0.031, sign test) (see Table 11 in Appendix C). However, this improvement comes with reduced search diversity, with the proportion of distinct method families across proposals decreasing from 52.5% to 40.0% on average during the optimisation (*p* = 0.001, sign test). Additionally, prediction calibration does not significantly improve, with the mean absolute error between predicted and observed performance only decreasing from 0.076 to 0.068 (*p* = 0.096, sign test). The optimisation traces for all 18 scenarios are provided in Figure 9 in Appendix C.

### 5.4 Analysis of model reasoning, size and the importance of the dossier

To better understand the optimisation process and the components driving performance gains, we conduct a series of ablation studies examining model reasoning, the contribution of each *dossier* section, and the effect of LLM size. We summarise our main findings below.

#### Citation frequency does not reflect section importance

First, we examine how the *proposer* agent uses the information in the *dossier*. Figure 4 shows the proportion of proposals that obeyed (i.e. hard constraints) or cited (i.e. used during candidate generation) each *dossier* section, alongside the effect on performance when sections are removed. We observe that the sections most frequently used by the agent are not necessarily those that contribute most to downstream performance. Further analysis over all 18 scenarios shows that the most frequently cited section of the *dossier* is the per-stage summary (§4), which appears in 81.5–87.0% of all proposals, compared with 11.2–46.6% for top-ranked pipelines (§2), 0–0.7% for rejected/retired pipelines (§3), and 1.5–10.9% for model predictions (§6) (see Table 12 in Appendix D). Yet removing §4 has little effect on performance (*−*0.006 *±* 0.005 AUC), whereas removing §2 substantially hurts performance (*−*0.032 *±* 0.008). In fact, dropping §2 is the only ablation to reduce performance in all four paired runs (see Table 13 in Appendix D), highlighting that the record of top-performing candidates is key for guiding the search

**Figure 4:**
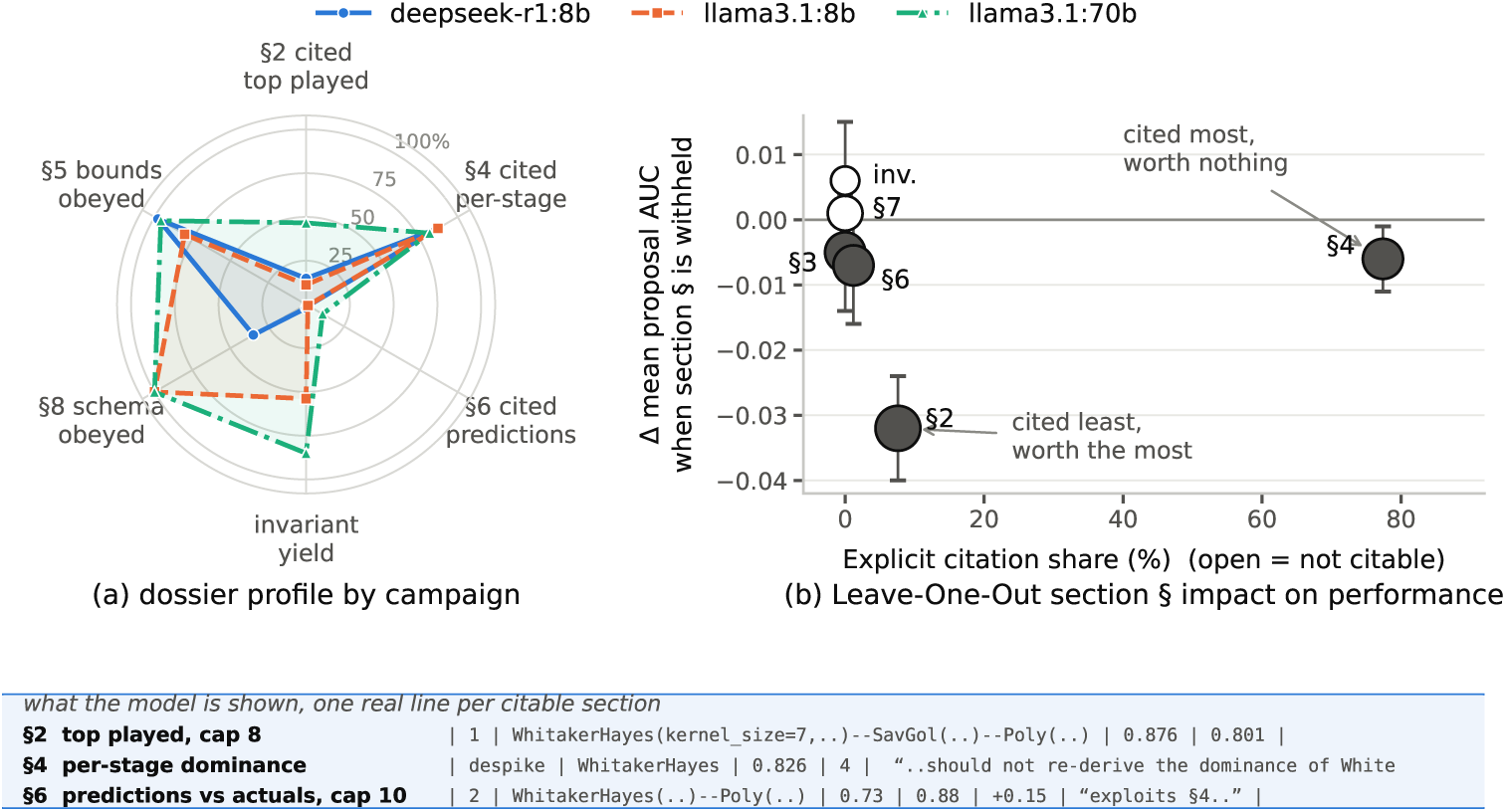
In-sample analysis of the reasoning of the *proposal* model and the importance of the *dossier*. (a) Proportion of proposals obeying/citing each *dossier* section. (b) Change in LLM AUC predictive performance when a *dossier* section is dropped.

#### The *dossier* is key for generating valid pipelines

We next examine which information in the *dossier* enables the generation of valid pipelines. We observe that when given only the task description, models return well-formed outputs only 3.2% of the time, and none of these is a valid pipeline (see Table 13 in Appendix D). Adding the output contract yields well-formed outputs, but 7,086 of the 7,140 proposed pipelines were still rejected as they contained unrecognised parameter names rather than out-of-range values. Adding section §5 resolved this issue, with only 274 rejections that were genuine constraint violations. This suggests that the registry supplies the API vocabulary required to generate valid pipelines, which, together with incorporating relevant parameter bounds during the optimisation process, substantially reduces the number of invalid parameters and thus pipelines explored. Removing individual dossier components generally reduced performance. Only removing §7 diagnostics or the self-model (§6 and invariants) yields a slightly higher mean proposal AUC (+0.001), well within the observed variability. The largest reductions occurred when removing §2 top-performing history (*−*0.020 *±* 0.010) or all optimisation history (*−*0.027 *±* 0.012), highlighting the importance of historical feedback. Disabling the *reflector* reduced mean proposal AUC by 0.009 *±* 0.005, with lower performance in 10 of 12 scenarios, yet the best achieved AUC across scenarios remained similar on average (0.723 versus 0.721; Table 13).

#### Increasing model size improves calibration, but not optimisation performance

The 70B `llama3.1` model outperformed its smaller counterpart (8B) only in 6 of 18 scenarios, with no significant difference in achieved AUC (mean difference *−*0.006 AUC, Wilcoxon *p* = 0.25). However, mean absolute prediction error decreases from 0.071 to 0.041 between 70B and 8B (*p* = 5.3 *×* 10*^−^*^5^, signed test), and the rank correlation between predicted and measured AUC goes from +0.507 to +0.791 (see Table 14 and Figure 10 in Appendix D). Prediction errors were concentrated in the baseline correction stage (see Table 15 in Appendix D), suggesting a systematic pipeline-specific bias rather than a model-specific effect. Thus, in our experiments, model scale improves candidate assessment but does not yield better pipelines within the search budget.

## 6 Conclusion

In conclusion, we have developed here an LLM-based agentic system for automated preprocessing optimisation in spectroscopy. We benchmarked our framework on 18 scenarios across six clinical datasets and three spectroscopic modalities. Our agentic optimisation achieved the highest out-of-sample AUC in four of the six datasets, while evaluating fewer than 5% as many pipelines as exhaustive grid search. We empirically show that our *dossier* infrastructure is a key contributor to performance, guiding pipeline generation, optimisation and search. Looking ahead, our system provides a proof of concept for more autonomous, reasoning-based workflows in spectroscopy. We look forward to seeing how advances in LLM reasoning, inference efficiency and agent orchestration could further improve both the predictive performance and computational efficiency of agentic preprocessing optimisation.

## AI use statement

In this work, we used generative AI tools to assist with code organisation, formatting, and debugging. We have not used generative AI tools to generate or modify the scientific methodology, experimental design, data, results, analysis, or conclusions. Additionally, generative AI tools were used to assist with code documentation and presentation. All AI-assisted code and content were reviewed, tested, and verified by the authors. We take full responsibility for the final content of this work, including all text, claims, code, results, and other artifacts produced with the aid of generative AI.

## Reproducibility statement

All datasets used in this study are publicly available and are listed below with their corresponding sources: (i) SERS dataset from Paraskevaidi et al. (2018) [https://figshare.com/articles/dataset/Raman_spectroscopic_techniques_to_detect_ovarian_cancer_biomarkers_in_blood_plasma/6744206]; (ii) ATR-FTIR (plasma) from Giamougiannis et al. (2021b) [https://figshare.com/articles/dataset/Raman_Ovarian_Cancer/14562312]; (iii) ATR-FTIR (serum) from Giamougiannis et al. (2021a) [https://figshare.com/articles/dataset/Raman_Ovarian_Cancer/14562312]; (iv) DRS dataset from Poh et al. (2020) [https://journals.plos.org/plosone/article?id=10.1371/journal.pone.0228923].

All experiments were performed using fixed random seeds to ensure reproducibility. Each experiment was repeated five times, using the same set of random seeds across all methods to ensure a consistent basis for comparison. Unless otherwise specified, data splits and all other experimental settings were kept identical across methods.

For experiments involving LLMs, we used the following models: (i) DeepSeek-R1:8B, (ii) Llama 3.1:8B, and (iii) Llama 3.1:70B, as described in Section 4. All models were run using the same experimental pipeline and evaluation protocol. Relevant parameters and inference settings are reported in the corresponding experimental sections.

Experiments were performed on the following hardware configurations: (i) workstation: AMD Ryzen Threadripper PRO 7975WX (up to 5.3 GHz), 256 GB DDR5 RAM, 4 TB M.2 storage in RAID 1, 24 TB HDD storage in RAID 1, and NVIDIA RTX PRO 6000 GPU; (ii) laptop: Apple MacBook Pro with M4 Max chip and 48 GB unified memory.

## Acknowledgments

G.P and D.G. acknowledge support from the UKRI Centre for Doctoral Training in AI for Healthcare (EP/S023283/1). D.G. acknowledges support from the EPSRC Postdoctoral Pathway scheme (EP/Z534870/1). D.G. and M.M.S. acknowledge funding from the Research Council of Norway through its Centres of Excellence scheme (262613). A.F.-G. acknowledges support from the Schmidt Science Fellows, in partnership with the Rhodes Trust. H.-C.N. and M.M.S. acknowledge support from the EPSRC Digital Health Hub for Antimicrobial Resistance (EP/X031276/1). H.-C.N. acknowledges support from the Medical Sciences Internal Fund: Pump priming funded by the University of Oxford. M.M.S. acknowledges support from the Royal Academy of Engineering Chair in Emerging Technologies award (CiET2021\94) and the University of Oxford Strategic Research Fund. M.B. acknowledges support from the Engineering and Physical Sciences Research Council (EP/N014529/1, funding the EPSRC Centre for Mathematics of Precision Healthcare at Imperial College London, and EP/T027258/1).

## A System architecture

**Figure 5:**
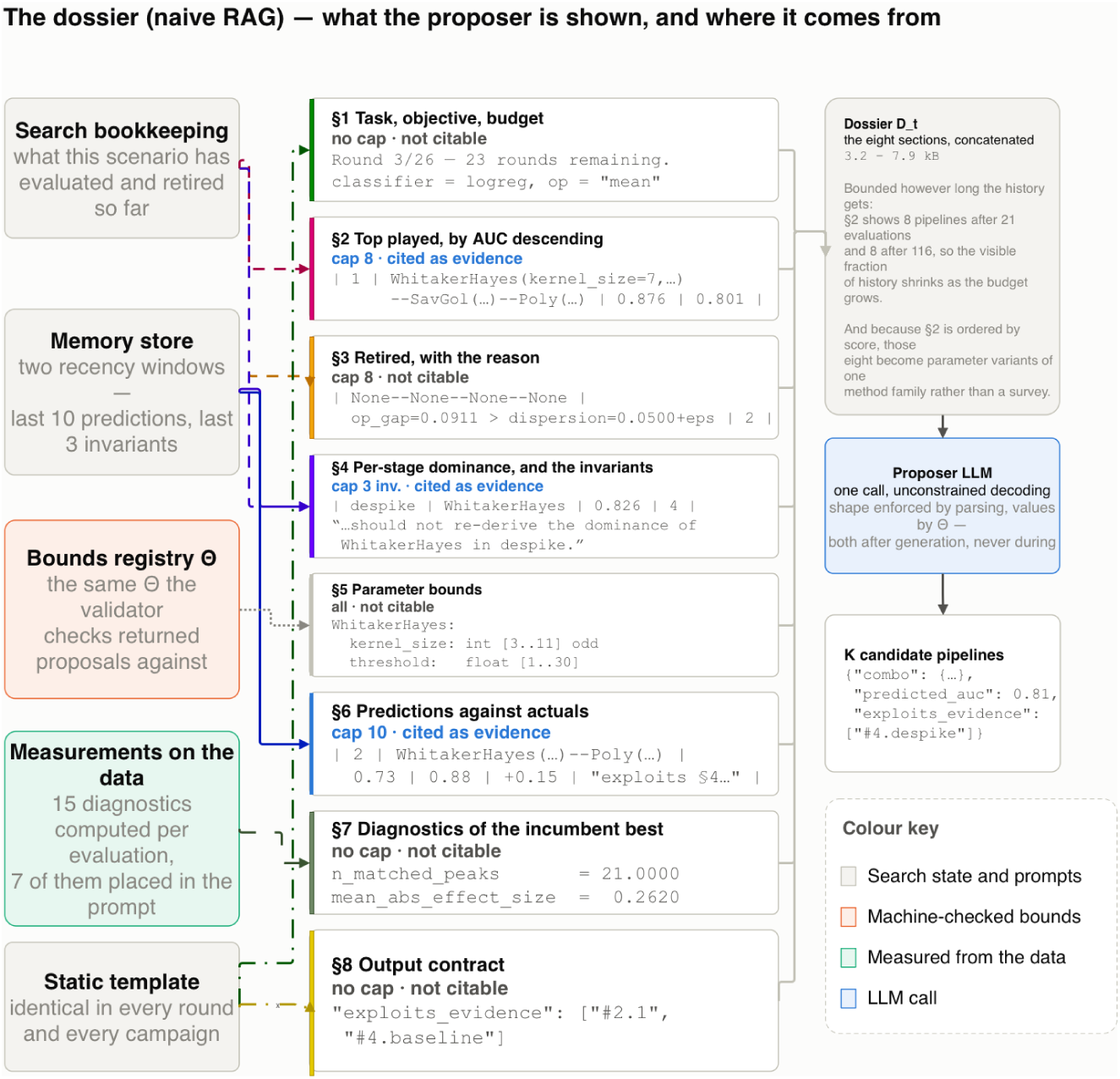
Anatomy of the dossier *D_t_*: the eight sections and the memory reads that populate them.

**Table 2:** Anatomy of the dossier. Cap is the number of entries retained. Cited is whether the content of the section has been explicitly cited by the Agent/LLM.

| § | Contents | Cap | Cited? | Example line |
|---|---|---|---|---|
| 1 | Task, objective, budget | — | no | Round 3/26 -- 23 rounds remaining.<br>classifier = logreg, op="mean" |
| 2 | Top played, by AUC desc | 8 | yes | \| 1 \| WhitakerHayes(kernel_size=7,...)-SavGol(...)-Poly(...) \| 0.876 \| 0.801 \| |
| 3 | Retired, with reason | 8 | no<sup>*</sup> | \| None-None-None-None \| op_gap=0.0911 > dispersion=0.0500+eps \| 2 \| |
| 4 | Per-stage dominance, plus reflector invariants | 3 inv. | yes | \| despike \| WhitakerHayes \| 0.826 \| 4 \|<br>“. . . should not re-derive the dominance of WhitakerHayes in despike.” |
| 5 | Parameter bounds | all | no | WhitakerHayes:<br>kernel_size: int [3..11] odd<br>threshold: float [1..30] |
| 6 | Predictions vs. actuals | 10 | yes | \| 2 \| WhitakerHayes(...)-Poly(...) \| 0.73 \| 0.88 \| +0.15 \| "exploits §4..." \| |
| 7 | Diagnostics of current best | — | no | n_matched_peaks = 21.0000<br>mean_abs_effect_size = 0.2620 |
| 8 | Output contract | — | no | "exploits_evidence": ["#2.1", "#4.baseline"] |

<sup>\*</sup> §3 appears in the rules only as a prohibition ("do not repeat a combo from §2 or §3"), never as citable evidence.

**Figure 6:**
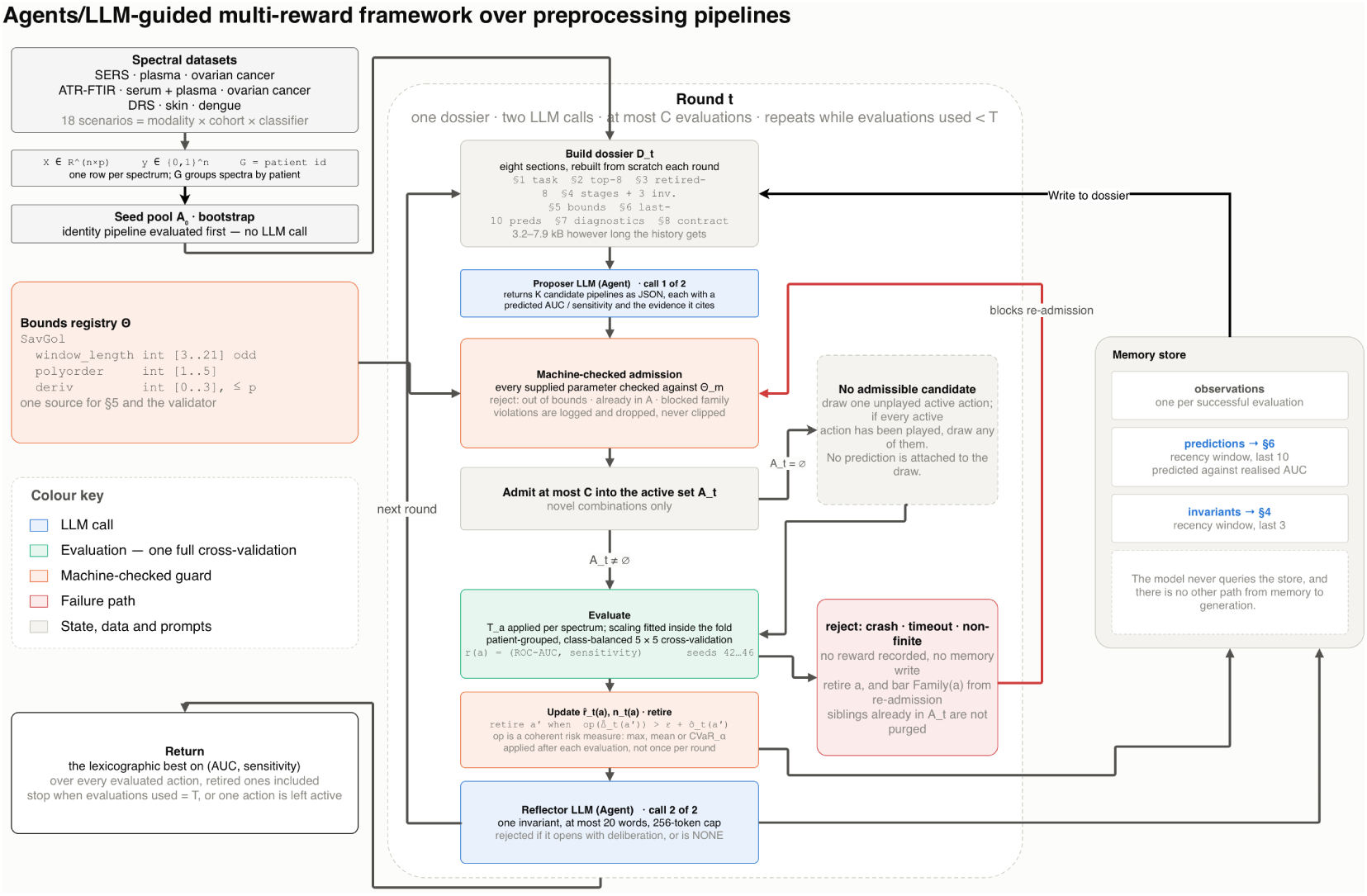
Architecture of the LLM-guided optimiser. The proposer, reflector, and memory store interact through the dossier rebuilt each round.

**Table 3:** The spectral diagnostics reported to the model in dossier §7, computed on the transformed matrix for the current best pipeline. During held-out evaluation, these diagnostics are computed exclusively on the training data available to the inner optimisation procedure, with no access to the outer test set. Example values are for `WhitakerHayes-SavGol-Poly` on the SERS ovarian-cancer task.

| Metric | What it measures | Example |
|---|---|---|
| *Class separability: did the transform help?* |  |  |
| mean_abs_effect_size | mean \|Cohen’s *d*\| over wavenumbers | 0.2620 |
| max_abs_effect_size | the single most discriminative channel | 1.3076 |
| mean_abs_auc_deviation | mean \|per-channel AUC − 0.5\| | 0.0768 |
| *Shape preservation: how much did it distort?* |  |  |
| pearson_mean_spectra | correlation of the two class-mean spectra | 0.9883 |
| cosine_mean_spectra | cosine similarity of the same pair | 0.9883 |
| *Peak structure: did bands survive, and move?* |  |  |
| n_matched_peaks | peaks present in both class means | 21 |
| mean_abs_peak_shift | mean \|Δposition\|, axis units | 0.5217 |

## B Datasets

**Table 4:**
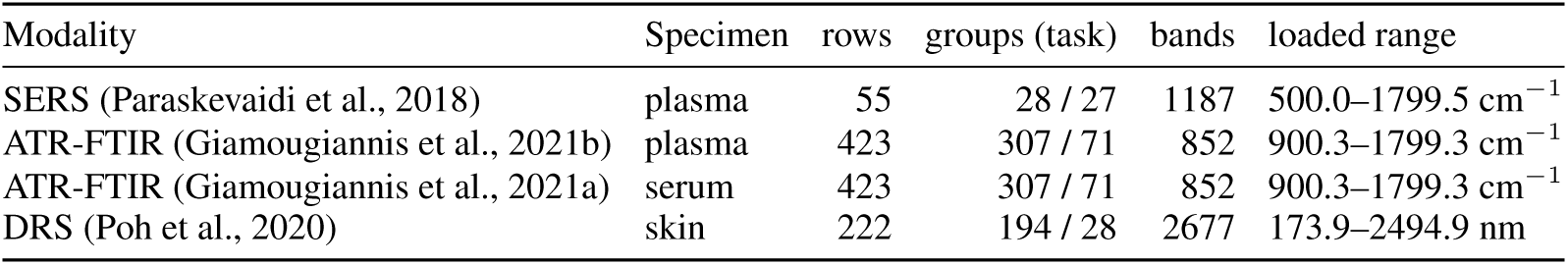
Extended information about the used datasets.

| Modality | Specimen | rows | groups (task) | bands | loaded range |
|---|---|---|---|---|---|
| SERS (Paraskevaidi et al., 2018) | plasma | 55 | 28 / 27 | 1187 | 500.0–1799.5 cm<sup>−1</sup> |
| ATR-FTIR (Giamougiannis et al., 2021b) | plasma | 423 | 307 / 71 | 852 | 900.3–1799.3 cm<sup>−1</sup> |
| ATR-FTIR (Giamougiannis et al., 2021a) | serum | 423 | 307 / 71 | 852 | 900.3–1799.3 cm<sup>−1</sup> |
| DRS (Poh et al., 2020) | skin | 222 | 194 / 28 | 2677 | 173.9–2494.9 nm |

## C EXTENDED RESULTS

**Figure 7:**
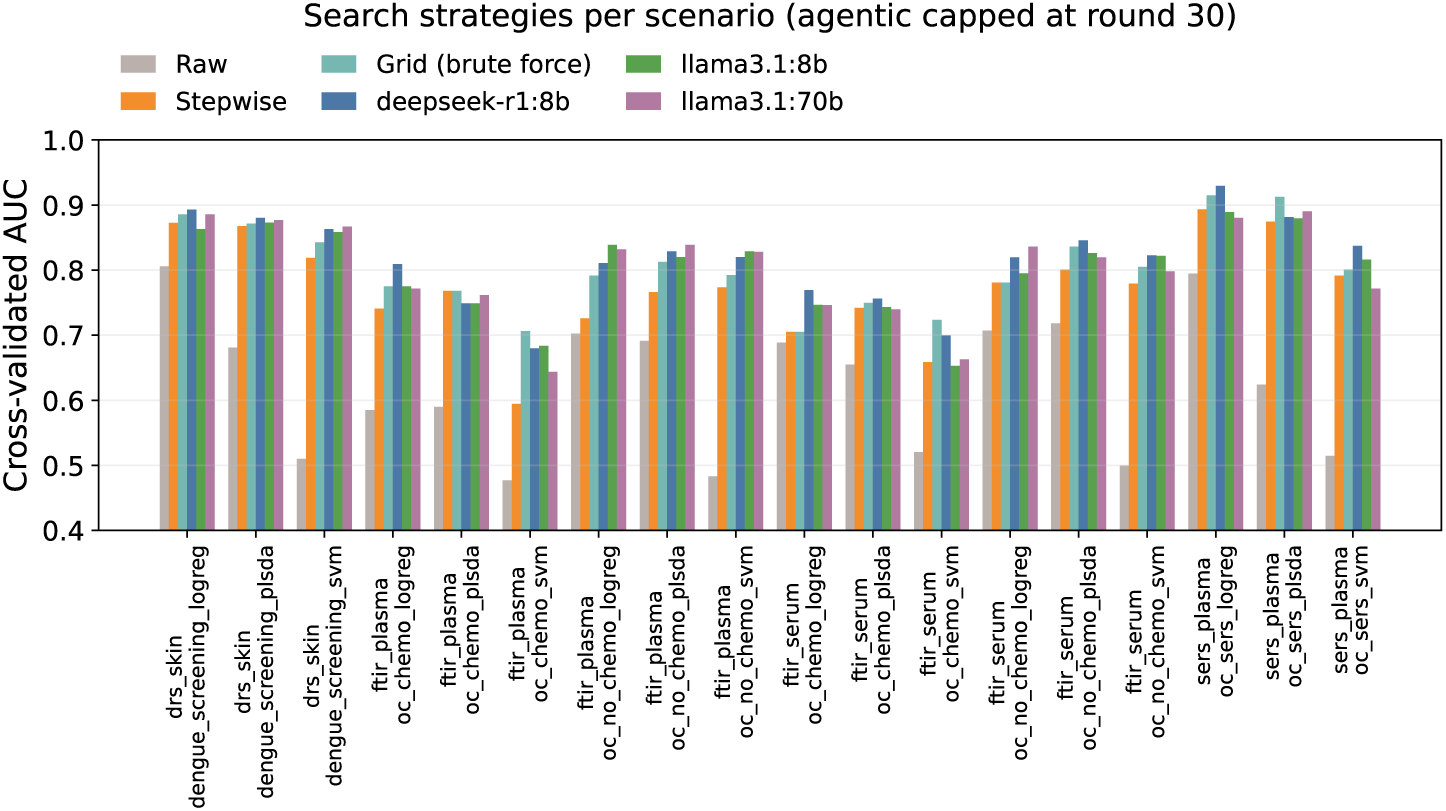
Benchmark results (30 rounds, in-sample). Top-1 classification performance achieved by each preprocessing optimisation strategy across the 18 scenarios.

**Figure 8:**
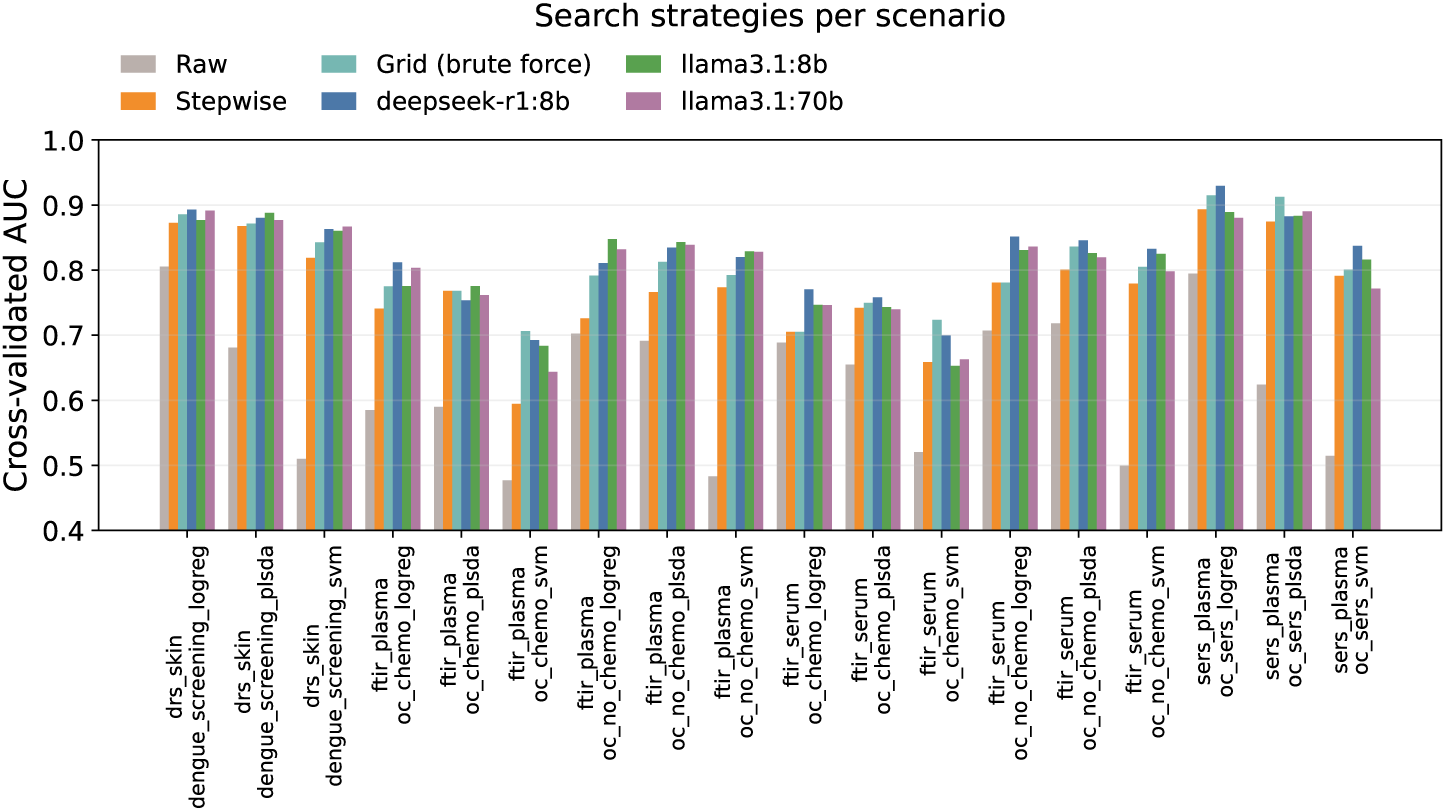
Extended benchmark results (120 evaluations, in-sample). Top-1 classification performance achieved by each preprocessing optimisation strategy across the 18 scenarios.

**Table 5:**
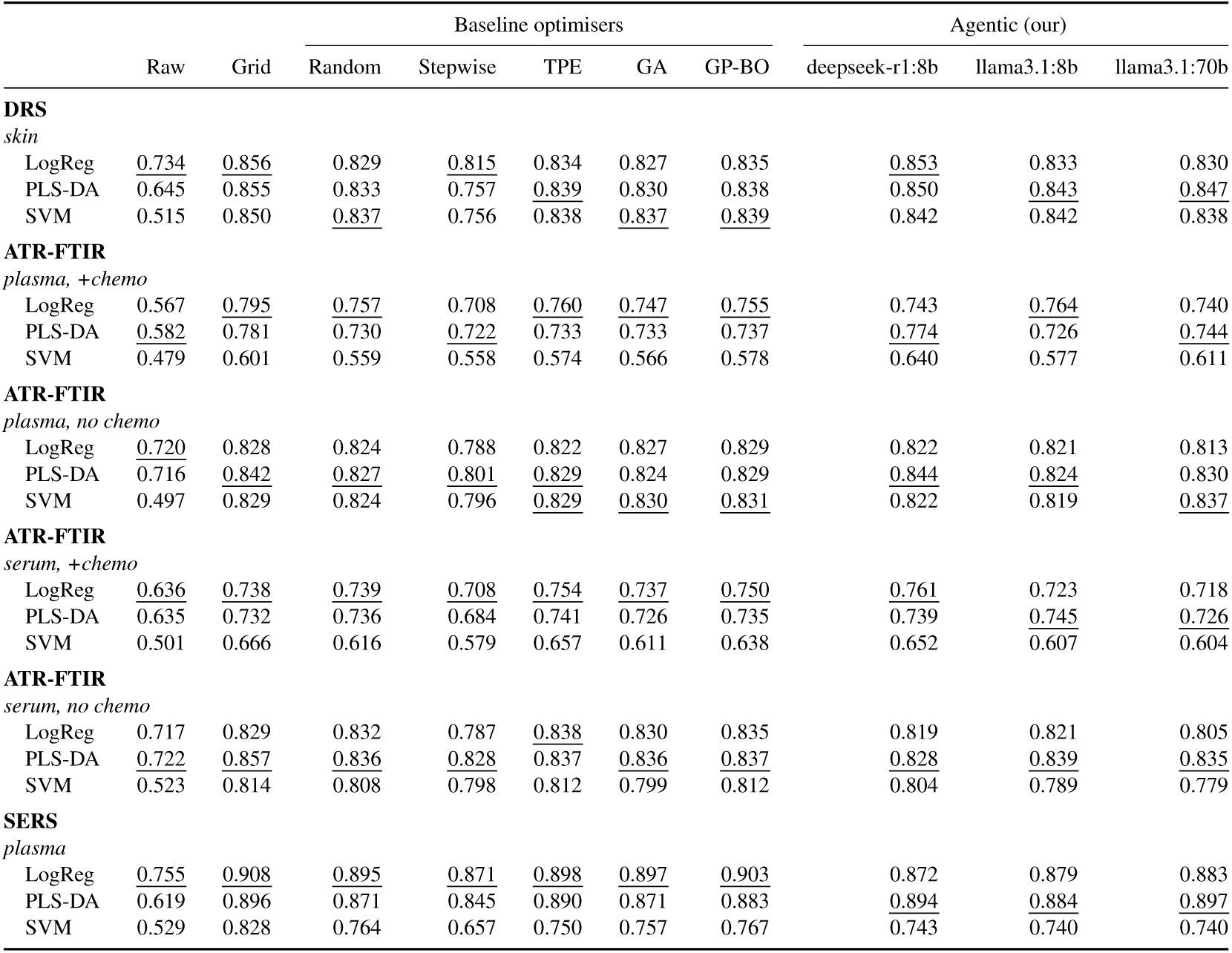
In-sample AUC scores (30 optimisation rounds). The underscored values show the best AUC for each method (raw, random, agentic, etc) and dataset. The classifier from each underlined combination was used to generate the out-of-sample Table 1.

|  |  |  | Baseline optimisers |  |  |  |  | Agentic (our) |  |  |
|---|---|---|---|---|---|---|---|---|---|---|
|  | Raw | Grid | Random | Stepwise | TPE | GA | GP-BO | deepseek-r1:8b | llama3.1:8b | llama3.1:70b |
| **DRS** |  |  |  |  |  |  |  |  |  |  |
| *skin* |  |  |  |  |  |  |  |  |  |  |
| LogReg | 0.734 | 0.856 | 0.829 | 0.815 | 0.834 | 0.827 | 0.835 | 0.853 | 0.833 | 0.830 |
| PLS-DA | 0.645 | 0.855 | 0.833 | 0.757 | 0.839 | 0.830 | 0.838 | 0.850 | 0.843 | 0.847 |
| SVM | 0.515 | 0.850 | 0.837 | 0.756 | 0.838 | 0.837 | 0.839 | 0.842 | 0.842 | 0.838 |
| **ATR-FTIR** |  |  |  |  |  |  |  |  |  |  |
| *plasma, +chemo* |  |  |  |  |  |  |  |  |  |  |
| LogReg | 0.567 | 0.795 | 0.757 | 0.708 | 0.760 | 0.747 | 0.755 | 0.743 | 0.764 | 0.740 |
| PLS-DA | 0.582 | 0.781 | 0.730 | 0.722 | 0.733 | 0.733 | 0.737 | 0.774 | 0.726 | 0.744 |
| SVM | 0.479 | 0.601 | 0.559 | 0.558 | 0.574 | 0.566 | 0.578 | 0.640 | 0.577 | 0.611 |
| **ATR-FTIR** |  |  |  |  |  |  |  |  |  |  |
| *plasma, no chemo* |  |  |  |  |  |  |  |  |  |  |
| LogReg | 0.720 | 0.828 | 0.824 | 0.788 | 0.822 | 0.827 | 0.829 | 0.822 | 0.821 | 0.813 |
| PLS-DA | 0.716 | 0.842 | 0.827 | 0.801 | 0.829 | 0.824 | 0.829 | 0.844 | 0.824 | 0.830 |
| SVM | 0.497 | 0.829 | 0.824 | 0.796 | 0.829 | 0.830 | 0.831 | 0.822 | 0.819 | 0.837 |
| **ATR-FTIR** |  |  |  |  |  |  |  |  |  |  |
| *serum, +chemo* |  |  |  |  |  |  |  |  |  |  |
| LogReg | 0.636 | 0.738 | 0.739 | 0.708 | 0.754 | 0.737 | 0.750 | 0.761 | 0.723 | 0.718 |
| PLS-DA | 0.635 | 0.732 | 0.736 | 0.684 | 0.741 | 0.726 | 0.735 | 0.739 | 0.745 | 0.726 |
| SVM | 0.501 | 0.666 | 0.616 | 0.579 | 0.657 | 0.611 | 0.638 | 0.652 | 0.607 | 0.604 |
| **ATR-FTIR** |  |  |  |  |  |  |  |  |  |  |
| *serum, no chemo* |  |  |  |  |  |  |  |  |  |  |
| LogReg | 0.717 | 0.829 | 0.832 | 0.787 | 0.838 | 0.830 | 0.835 | 0.819 | 0.821 | 0.805 |
| PLS-DA | 0.722 | 0.857 | 0.836 | 0.828 | 0.837 | 0.836 | 0.837 | 0.828 | 0.839 | 0.835 |
| SVM | 0.523 | 0.814 | 0.808 | 0.798 | 0.812 | 0.799 | 0.812 | 0.804 | 0.789 | 0.779 |
| **SERS** |  |  |  |  |  |  |  |  |  |  |
| *plasma* |  |  |  |  |  |  |  |  |  |  |
| LogReg | 0.755 | 0.908 | 0.895 | 0.871 | 0.898 | 0.897 | 0.903 | 0.872 | 0.879 | 0.883 |
| PLS-DA | 0.619 | 0.896 | 0.871 | 0.845 | 0.890 | 0.871 | 0.883 | 0.894 | 0.884 | 0.897 |
| SVM | 0.529 | 0.828 | 0.764 | 0.657 | 0.750 | 0.757 | 0.767 | 0.743 | 0.740 | 0.740 |

**Table 6:** Out-of-sample AUC scores (30 optimisation rounds). The best AUC values for each scenario are marked in bold.

|  |  |  | Baseline optimisers |  |  |  |  | Agentic (our) |  |  |
|---|---|---|---|---|---|---|---|---|---|---|
|  | Raw | Grid | Random | Stepwise | TPE | GA | GP-BO | DeepSeek-R1<br>(8B) | Llama 3.1<br>(8B) | Llama 3.1<br>(70B) |
| **DRS** |  |  |  |  |  |  |  |  |  |  |
| *skin* |  |  |  |  |  |  |  |  |  |  |
| LogReg | 0.675 | 0.739 | 0.745 | 0.729 | 0.752 | 0.740 | 0.744 | 0.793 | **0.803** | 0.761 |
| PLS-DA | 0.656 | 0.784 | 0.786 | 0.752 | 0.787 | 0.815 | 0.788 | 0.838 | 0.838 | **0.840** |
| SVM | 0.656 | 0.767 | 0.765 | 0.660 | 0.773 | 0.784 | 0.772 | 0.771 | 0.776 | **0.840** |
| **ATR-FTIR** |  |  |  |  |  |  |  |  |  |  |
| *plasma, +chemo* |  |  |  |  |  |  |  |  |  |  |
| LogReg | 0.647 | 0.719 | 0.777 | 0.739 | 0.746 | 0.748 | 0.748 | 0.703 | 0.722 | **0.796** |
| PLS-DA | 0.614 | 0.702 | 0.744 | 0.753 | 0.737 | 0.707 | 0.734 | 0.741 | **0.760** | 0.743 |
| SVM | 0.545 | 0.531 | 0.508 | 0.542 | 0.566 | 0.535 | 0.524 | 0.595 | **0.637** | 0.556 |
| **ATR-FTIR** |  |  |  |  |  |  |  |  |  |  |
| *plasma, no chemo* |  |  |  |  |  |  |  |  |  |  |
| LogReg | 0.681 | 0.727 | 0.718 | 0.711 | 0.741 | 0.719 | 0.721 | 0.771 | 0.752 | **0.788** |
| PLS-DA | 0.693 | 0.772 | 0.757 | 0.753 | 0.757 | 0.770 | 0.765 | 0.773 | **0.785** | 0.781 |
| SVM | 0.426 | 0.724 | 0.742 | **0.764** | 0.755 | 0.737 | 0.750 | 0.748 | 0.761 | 0.723 |
| **ATR-FTIR** |  |  |  |  |  |  |  |  |  |  |
| *serum, +chemo* |  |  |  |  |  |  |  |  |  |  |
| LogReg | 0.676 | 0.675 | 0.689 | 0.684 | 0.718 | 0.703 | 0.704 | 0.699 | 0.733 | **0.747** |
| PLS-DA | 0.690 | 0.679 | 0.685 | 0.687 | 0.705 | 0.700 | 0.702 | 0.696 | 0.696 | **0.743** |
| SVM | 0.531 | 0.607 | 0.587 | 0.579 | 0.607 | 0.560 | 0.573 | 0.589 | **0.621** | 0.595 |
| **ATR-FTIR** |  |  |  |  |  |  |  |  |  |  |
| *serum, no chemo* |  |  |  |  |  |  |  |  |  |  |
| LogReg | 0.693 | 0.796 | 0.786 | 0.797 | 0.803 | 0.800 | 0.805 | 0.810 | **0.845** | 0.739 |
| PLS-DA | 0.699 | 0.759 | 0.802 | **0.827** | 0.794 | 0.795 | 0.799 | 0.779 | 0.815 | 0.753 |
| SVM | 0.529 | 0.801 | 0.800 | 0.802 | **0.814** | 0.803 | 0.808 | 0.804 | 0.795 | 0.812 |
| **SERS** |  |  |  |  |  |  |  |  |  |  |
| *plasma* |  |  |  |  |  |  |  |  |  |  |
| LogReg | 0.717 | 0.800 | 0.840 | 0.850 | 0.837 | 0.842 | 0.818 | 0.867 | 0.733 | **0.900** |
| PLS-DA | 0.725 | 0.867 | 0.788 | 0.783 | 0.778 | 0.820 | 0.837 | **0.917** | 0.867 | 0.867 |
| SVM | 0.500 | 0.617 | **0.668** | 0.633 | 0.625 | 0.620 | 0.622 | 0.517 | 0.567 | 0.567 |

**Table 7:**
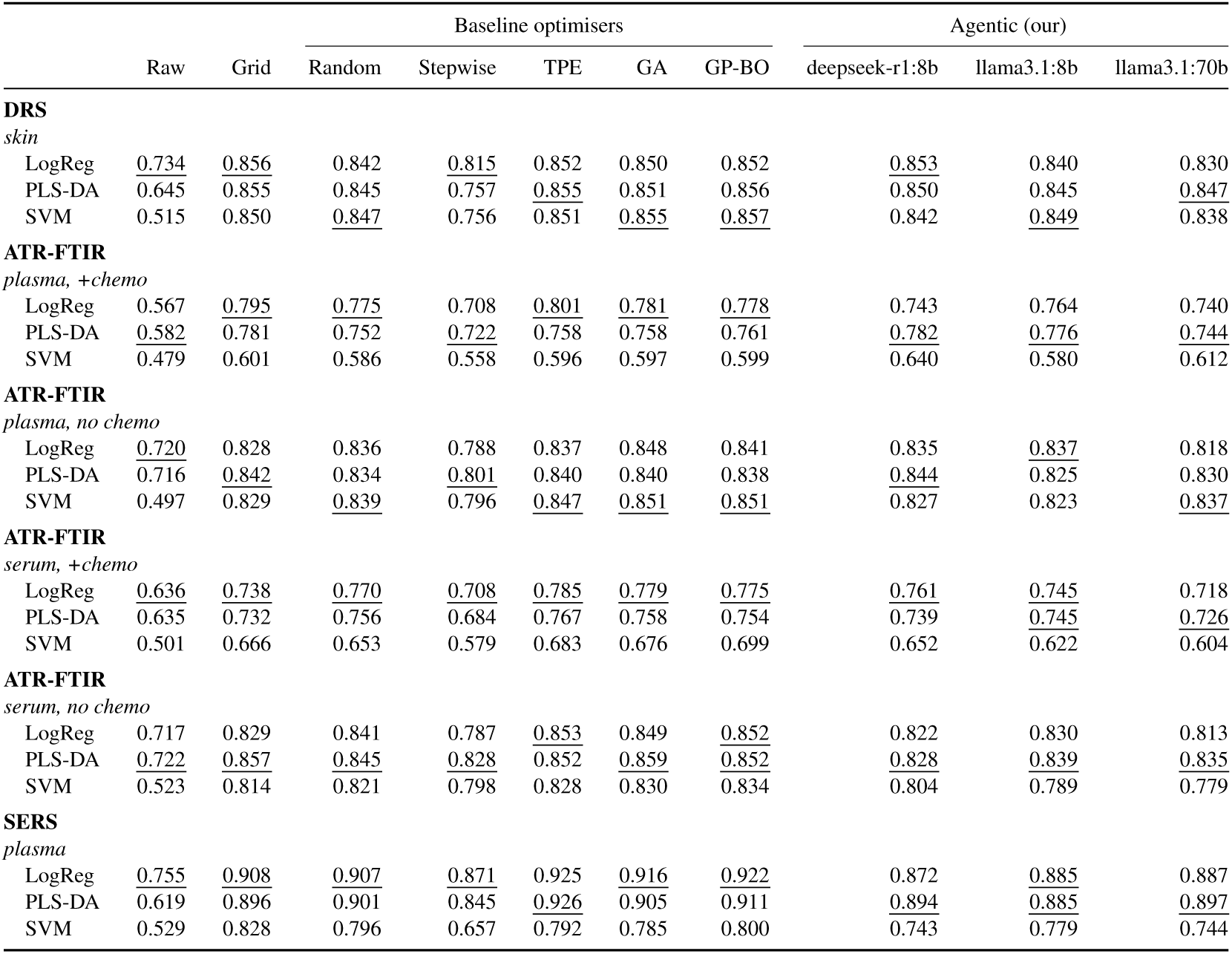
In-sample AUC scores (120 evaluations). The underscored values show the best AUC for each method (raw, random, agentic, etc) and dataset. Baseline optimisation methods are capped at 120 rounds, 1 pipeline evaluation per round. The agentic framework evaluates a maximum of 5 pipelines per round.

|  |  |  | Baseline optimisers |  |  |  |  | Agentic (our) |  |  |
|---|---|---|---|---|---|---|---|---|---|---|
|  | Raw | Grid | Random | Stepwise | TPE | GA | GP-BO | deepseek-r1:8b | llama3.1:8b | llama3.1:70b |
| **DRS** |  |  |  |  |  |  |  |  |  |  |
| *skin* |  |  |  |  |  |  |  |  |  |  |
| LogReg | 0.734 | 0.856 | 0.842 | 0.815 | 0.852 | 0.850 | 0.852 | 0.853 | 0.840 | 0.830 |
| PLS-DA | 0.645 | 0.855 | 0.845 | 0.757 | 0.855 | 0.851 | 0.856 | 0.850 | 0.845 | 0.847 |
| SVM | 0.515 | 0.850 | 0.847 | 0.756 | 0.851 | 0.855 | 0.857 | 0.842 | 0.849 | 0.838 |
| **ATR-FTIR** |  |  |  |  |  |  |  |  |  |  |
| *plasma, +chemo* |  |  |  |  |  |  |  |  |  |  |
| LogReg | 0.567 | 0.795 | 0.775 | 0.708 | 0.801 | 0.781 | 0.778 | 0.743 | 0.764 | 0.740 |
| PLS-DA | 0.582 | 0.781 | 0.752 | 0.722 | 0.758 | 0.758 | 0.761 | 0.782 | 0.776 | 0.744 |
| SVM | 0.479 | 0.601 | 0.586 | 0.558 | 0.596 | 0.597 | 0.599 | 0.640 | 0.580 | 0.612 |
| **ATR-FTIR** |  |  |  |  |  |  |  |  |  |  |
| *plasma, no chemo* |  |  |  |  |  |  |  |  |  |  |
| LogReg | 0.720 | 0.828 | 0.836 | 0.788 | 0.837 | 0.848 | 0.841 | 0.835 | 0.837 | 0.818 |
| PLS-DA | 0.716 | 0.842 | 0.834 | 0.801 | 0.840 | 0.840 | 0.838 | 0.844 | 0.825 | 0.830 |
| SVM | 0.497 | 0.829 | 0.839 | 0.796 | 0.847 | 0.851 | 0.851 | 0.827 | 0.823 | 0.837 |
| **ATR-FTIR** |  |  |  |  |  |  |  |  |  |  |
| *serum, +chemo* |  |  |  |  |  |  |  |  |  |  |
| LogReg | 0.636 | 0.738 | 0.770 | 0.708 | 0.785 | 0.779 | 0.775 | 0.761 | 0.745 | 0.718 |
| PLS-DA | 0.635 | 0.732 | 0.756 | 0.684 | 0.767 | 0.758 | 0.754 | 0.739 | 0.745 | 0.726 |
| SVM | 0.501 | 0.666 | 0.653 | 0.579 | 0.683 | 0.676 | 0.699 | 0.652 | 0.622 | 0.604 |
| **ATR-FTIR** |  |  |  |  |  |  |  |  |  |  |
| *serum, no chemo* |  |  |  |  |  |  |  |  |  |  |
| LogReg | 0.717 | 0.829 | 0.841 | 0.787 | 0.853 | 0.849 | 0.852 | 0.822 | 0.830 | 0.813 |
| PLS-DA | 0.722 | 0.857 | 0.845 | 0.828 | 0.852 | 0.859 | 0.852 | 0.828 | 0.839 | 0.835 |
| SVM | 0.523 | 0.814 | 0.821 | 0.798 | 0.828 | 0.830 | 0.834 | 0.804 | 0.789 | 0.779 |
| **SERS** |  |  |  |  |  |  |  |  |  |  |
| *plasma* |  |  |  |  |  |  |  |  |  |  |
| LogReg | 0.755 | 0.908 | 0.907 | 0.871 | 0.925 | 0.916 | 0.922 | 0.872 | 0.885 | 0.887 |
| PLS-DA | 0.619 | 0.896 | 0.901 | 0.845 | 0.926 | 0.905 | 0.911 | 0.894 | 0.885 | 0.897 |
| SVM | 0.529 | 0.828 | 0.796 | 0.657 | 0.792 | 0.785 | 0.800 | 0.743 | 0.779 | 0.744 |

**Table 8:**
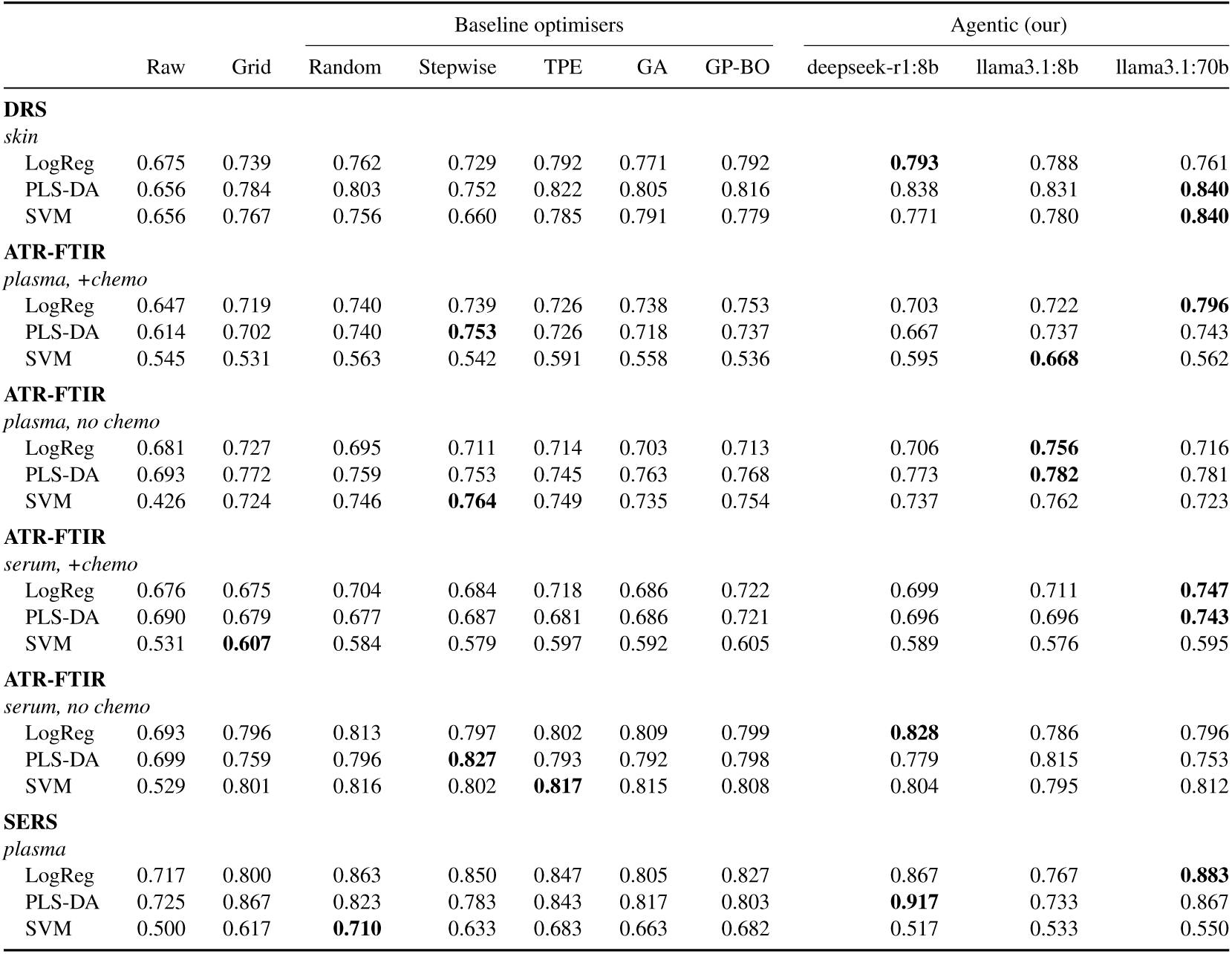
Out-of-sample AUC scores (120 evaluations). The best AUC values for each scenario are marked in bold. Baseline optimisation methods are capped at 120 rounds, 1 pipeline evaluation per round. The agentic framework evaluates a maximum of 5 pipelines per round.

|  |  |  | Baseline optimisers |  |  |  |  | Agentic (our) |  |  |
|---|---|---|---|---|---|---|---|---|---|---|
|  | Raw | Grid | Random | Stepwise | TPE | GA | GP-BO | deepseek-r1:8b | llama3.1:8b | llama3.1:70b |
| **DRS** |  |  |  |  |  |  |  |  |  |  |
| *skin* |  |  |  |  |  |  |  |  |  |  |
| LogReg | 0.675 | 0.739 | 0.762 | 0.729 | 0.792 | 0.771 | 0.792 | **0.793** | 0.788 | 0.761 |
| PLS-DA | 0.656 | 0.784 | 0.803 | 0.752 | 0.822 | 0.805 | 0.816 | 0.838 | 0.831 | **0.840** |
| SVM | 0.656 | 0.767 | 0.756 | 0.660 | 0.785 | 0.791 | 0.779 | 0.771 | 0.780 | **0.840** |
| **ATR-FTIR** |  |  |  |  |  |  |  |  |  |  |
| *plasma, +chemo* |  |  |  |  |  |  |  |  |  |  |
| LogReg | 0.647 | 0.719 | 0.740 | 0.739 | 0.726 | 0.738 | 0.753 | 0.703 | 0.722 | **0.796** |
| PLS-DA | 0.614 | 0.702 | 0.740 | **0.753** | 0.726 | 0.718 | 0.737 | 0.667 | 0.737 | 0.743 |
| SVM | 0.545 | 0.531 | 0.563 | 0.542 | 0.591 | 0.558 | 0.536 | 0.595 | **0.668** | 0.562 |
| **ATR-FTIR** |  |  |  |  |  |  |  |  |  |  |
| *plasma, no chemo* |  |  |  |  |  |  |  |  |  |  |
| LogReg | 0.681 | 0.727 | 0.695 | 0.711 | 0.714 | 0.703 | 0.713 | 0.706 | **0.756** | 0.716 |
| PLS-DA | 0.693 | 0.772 | 0.759 | 0.753 | 0.745 | 0.763 | 0.768 | 0.773 | **0.782** | 0.781 |
| SVM | 0.426 | 0.724 | 0.746 | **0.764** | 0.749 | 0.735 | 0.754 | 0.737 | 0.762 | 0.723 |
| **ATR-FTIR** |  |  |  |  |  |  |  |  |  |  |
| *serum, +chemo* |  |  |  |  |  |  |  |  |  |  |
| LogReg | 0.676 | 0.675 | 0.704 | 0.684 | 0.718 | 0.686 | 0.722 | 0.699 | 0.711 | **0.747** |
| PLS-DA | 0.690 | 0.679 | 0.677 | 0.687 | 0.681 | 0.686 | 0.721 | 0.696 | 0.696 | **0.743** |
| SVM | 0.531 | **0.607** | 0.584 | 0.579 | 0.597 | 0.592 | 0.605 | 0.589 | 0.576 | 0.595 |
| **ATR-FTIR** |  |  |  |  |  |  |  |  |  |  |
| *serum, no chemo* |  |  |  |  |  |  |  |  |  |  |
| LogReg | 0.693 | 0.796 | 0.813 | 0.797 | 0.802 | 0.809 | 0.799 | **0.828** | 0.786 | 0.796 |
| PLS-DA | 0.699 | 0.759 | 0.796 | **0.827** | 0.793 | 0.792 | 0.798 | 0.779 | 0.815 | 0.753 |
| SVM | 0.529 | 0.801 | 0.816 | 0.802 | **0.817** | 0.815 | 0.808 | 0.804 | 0.795 | 0.812 |
| **SERS** |  |  |  |  |  |  |  |  |  |  |
| *plasma* |  |  |  |  |  |  |  |  |  |  |
| LogReg | 0.717 | 0.800 | 0.863 | 0.850 | 0.847 | 0.805 | 0.827 | 0.867 | 0.767 | **0.883** |
| PLS-DA | 0.725 | 0.867 | 0.823 | 0.783 | 0.843 | 0.817 | 0.803 | **0.917** | 0.733 | 0.867 |
| SVM | 0.500 | 0.617 | **0.710** | 0.633 | 0.683 | 0.663 | 0.682 | 0.517 | 0.533 | 0.550 |

**Table 9:** Wall-clock cost of all methods tested. Running times were measured on a workstation using 1*×* RTX Pro 6000 GPU and 15 cores from AMD Ryzen Threadripper PRO 7975WX CPU. The LLM & orchestration stage for agents includes building and updating the dossier, including the metrics, agent proposals and evaluation of the dossier results. Running times were recorded on a single repeat.

| Method | rounds | evals | wall clock (h) |  | per unit |  |
|---|---|---|---|---|---|---|
|  |  |  | total | LLM & orchestration | per eval (s) | per scen. (h) |
| Raw | — | 18 | <1min | — | 1 | <1min |
| Grid | — | 43560 | 70 | — | 5.8 | 3.9 |
| Random | — | 2160 | 1.74 | — | 2.9 | 0.1 |
| Stepwise | — | 635 | 0.5 | — | 3 | 0.03 |
| TPE | — | 2160 | 2.04 | — | 3.4 | 0.1 |
| GA | — | 2160 | 1.86 | — | 3.1 | 0.1 |
| GP-BO | — | 2160 | 1.98 | — | 3.3 | 0.1 |
| deepseek-r1:8b | 570 | 2160 | 7.53 | 6.15 | 12.5 | 0.4 |
| llama3.1:8b | 923 | 2160 | 9.51 | 7.94 | 15.9 | 0.5 |
| llama3.1:70b<sup>*</sup> | 640 | 2160 | 1.31 | 10.60 | 19.8 | 0.7 |

**Table 10:**
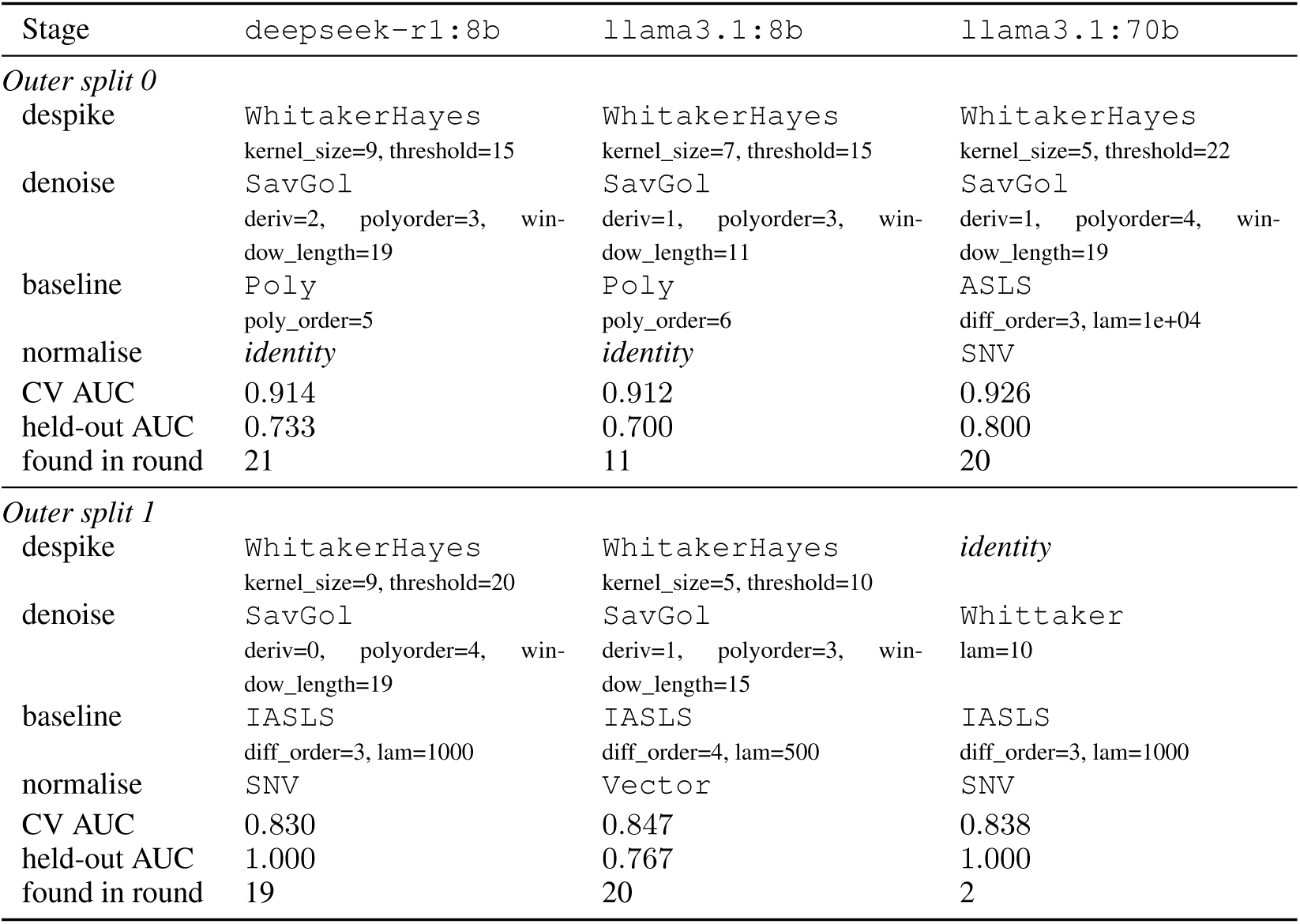
The highest-AUC pipeline each campaign found on sers_plasma_oc_sers_logreg, by stage, with the parameters as proposed. All three settle on a polynomial baseline and differ on everything else, including whether to despike or normalise at all.

| Stage | deepseek-r1:8b | llama3.1:8b | llama3.1:70b |
|---|---|---|---|
| *Outer split 0* |  |  |  |
| despike | WhitakerHayes<br>kernel_size=9, threshold=15 | WhitakerHayes<br>kernel_size=7, threshold=15 | WhitakerHayes<br>kernel_size=5, threshold=22 |
| denoise | SavGol<br>deriv=2, polyorder=3, win-dow_length=19 | SavGol<br>deriv=1, polyorder=3, win-dow_length=11 | SavGol<br>deriv=1, polyorder=4, win-dow_length=19 |
| baseline | Poly<br>poly_order=5 | Poly<br>poly_order=6 | ASLS<br>diff_order=3, lam=1e+04 |
| normalise | *identity* | *identity* | SNV |
| CV AUC | 0.914 | 0.912 | 0.926 |
| held-out AUC | 0.733 | 0.700 | 0.800 |
| found in round | 21 | 11 | 20 |
| *Outer split 1* |  |  |  |
| despike | WhitakerHayes<br>kernel_size=9, threshold=20 | WhitakerHayes<br>kernel_size=5, threshold=10 | *identity* |
| denoise | SavGol<br>deriv=0, polyorder=4, win-dow_length=19 | SavGol<br>deriv=1, polyorder=3, win-dow_length=15 | Whittaker<br>lam=10 |
| baseline | IASLS<br>diff_order=3, lam=1000 | IASLS<br>diff_order=4, lam=500 | IASLS<br>diff_order=3, lam=1000 |
| normalise | SNV | Vector | SNV |
| CV AUC | 0.830 | 0.847 | 0.838 |
| held-out AUC | 1.000 | 0.767 | 1.000 |
| found in round | 19 | 20 | 2 |

**Table 11:** Proposal quality and calibration across rounds for the `llama3.1:8b` campaign, comparing the first and last third of each scenario’s rounds over LLM-proposed pipelines only. *Scenarios improving* counts increases, except for prediction error where a decrease is the improvement. *p*-values are sign tests over the 18 scenarios.

| Measure | first third | last third | scenarios improving |
|---|---|---|---|
| mean AUC of proposals | 0.686 | **0.720** | 16/18 (*p* = 0.001) |
| hit rate above early baseline | 49.0% | **61.8%** | 14/18 (*p* = 0.031) |
| *ρ*(round, AUC) > 0 | — | — | 14/18 (*p* = 0.031) |
| mean \|prediction error\| | 0.076 | **0.068** | 13/18 (*p* = 0.096, n.s.) |
| distinct family combinations | **52.5%** | 40.0% | 2/18 (*p* = 0.001) |

**Figure 9:**
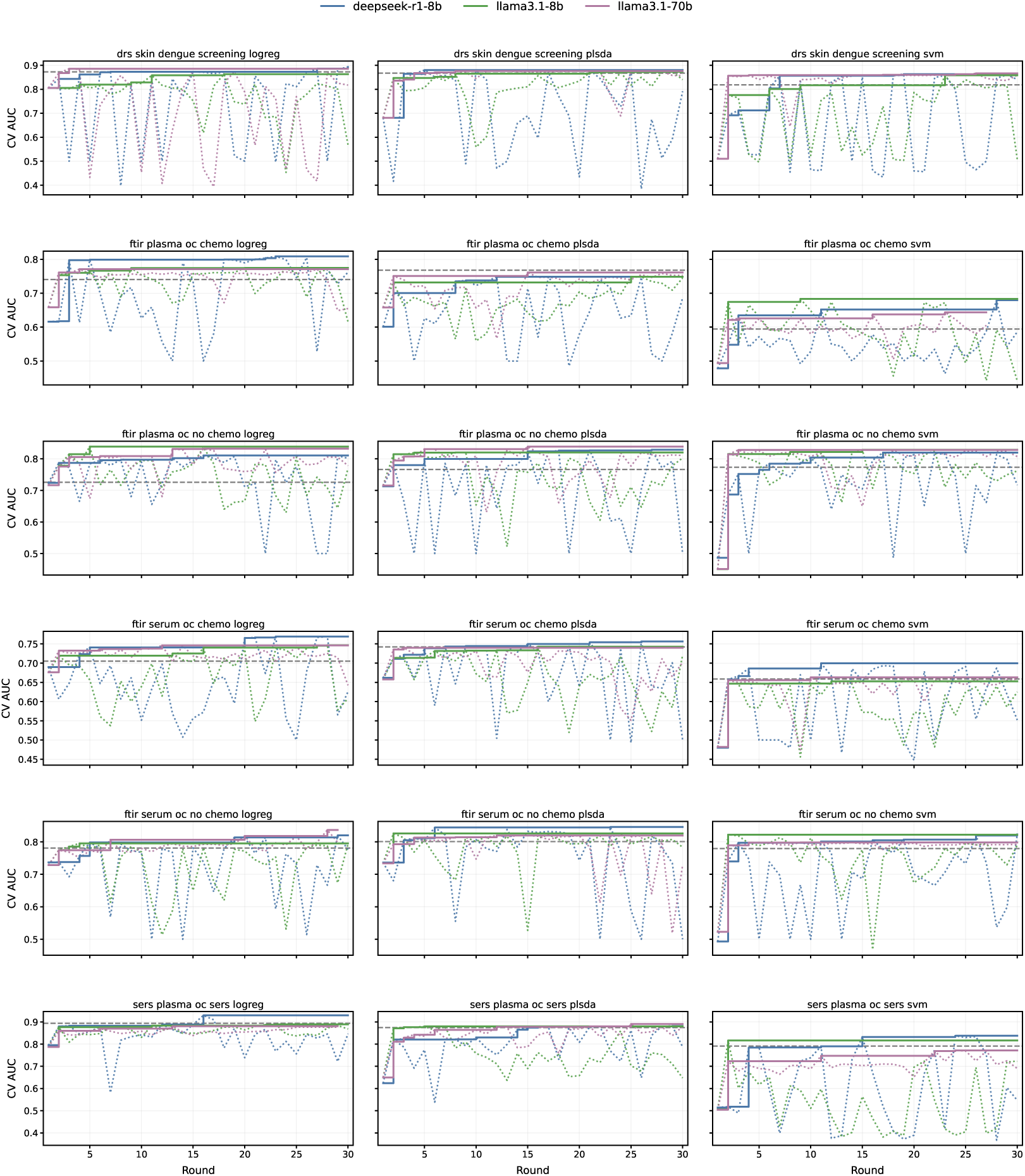
Figure 3 for every scenario. Rows are the (modality, specimen, task) group, columns the classifier, so the three panels of a row share a task and a *y*-scale. Dotted is the best of each round, the solid step the running best, the grey dashed line greedy stepwise. The pattern Section 5.2 draws from one scenario is general. The step lines flatten early almost everywhere, and the campaign with the highest step is often not the one whose dotted line sits closest to it.

## D Ablation studies

**Table 12:** Share of proposals citing each dossier section, pooled over all 18 scenarios. A section counts at most once per proposal. Only §2, §4 and §6 are presented as citable by the output contract, so §1, §5, §7 and §8 cannot be cited and are omitted. §3 is shown because it appears in the dossier’s rules, though only in a prohibition. The last three columns are observed rather than self-reported: bounds and schema compliance, and the share of rounds in which the reflector’s invariant survived the filters of Section 3.2. These are the numbers behind Figure 4(a).

|  |  | cited (self-reported) |  |  |  | obeyed (observed) |  |  |
|---|---|---|---|---|---|---|---|---|
| Campaign | proposals | §2 | §3 | §4 | §6 | §5 bounds | §8 schema | invariant |
| deepseek-r1:8b | 3216 | 14.9% | 0.0% | 86.9% | 1.5% | 97.8% | 34.8% | 2.0% |
| llama3.1:8b | 7384 | 11.2% | 0.0% | 87.0% | 1.2% | 80.1% | 100.0% | 53.8% |
| llama3.1:70b | 5758 | 46.6% | 0.7% | 81.5% | 10.9% | 95.8% | 100.0% | 85.0% |

**Table 13:** Ablation of dossier components and the reflector across 12 optimisation runs (3 models, 2 scenarios, 2 seeds). *Worse in* counts the runs in which an ablation underperformed the full configuration. The upper block removes individual dossier components. The lower block examines grouped removals and a progressively expanded control ladder (§1 *⊂* §1§8 *⊂* §1§5§8 *⊂* full). The bottom two ladder configurations admitted no proposals, so their AUC values reflect fallback draws rather than agent proposals. *^†^* indicates responses that could not be parsed. The reflector ablation disables the second agent to assess its contribution to proposal quality.

| Dossier | mean prop. AUC | Δ vs full | worse in | best AUC | §5 bounds | §8 contract |
|---|---|---|---|---|---|---|
| full dossier | 0.649 | — | — | 0.721 | 88.7% | 100.0% |
| – §2 top played | 0.630 | −0.020 ± 0.010 | 9/12 | 0.713 | 87.3% | 100.0% |
| – §3 eliminated | 0.643 | −0.006 ± 0.004 | 8/12 | 0.713 | 91.4% | 99.4% |
| – §4 dominance | 0.645 | −0.004 ± 0.002 | 8/12 | 0.721 | 92.8% | 100.0% |
| – §6 predictions | 0.647 | −0.002 ± 0.003 | 6/12 | 0.716 | 92.9% | 97.1% |
| – §7 diagnostics | 0.650 | +0.001 ± 0.004 | 7/12 | 0.726 | 90.0% | 97.0% |
| – invariants | 0.635 | −0.014 ± 0.005 | 10/12 | 0.714 | 93.2% | 100.0% |
| – all history (§2§4§7 + inv.) | 0.622 | −0.027 ± 0.012 | 9/12 | 0.713 | 91.0% | 93.0% |
| – self-model (§6 + inv.) | 0.650 | +0.001 ± 0.004 | 4/12 | 0.718 | 91.7% | 98.2% |
| controls only (§1§5§8) | 0.621 | −0.028 ± 0.005 | 12/12 | 0.718 | 85.9% | 100.0% |
| task + contract (§1§8) | — | — | — | — | 0.5% | 99.2% |
| task only (§1)<sup>†</sup> | — | — | — | — | 0.0% | 3.2% |
| – reflector (call off) | 0.640 | −0.009 ± 0.005 | 10/12 | 0.723 | 93.5% | 100.0% |

**Table 14:** Predicted against measured AUC, pooled over every proposal a campaign made. *Signed error* is measured minus predicted, so a negative value is over-prediction. Spearman is the rank correlation between predicted and measured AUC, with *^∗∗∗^* denoting *p <* 10*^−^*^3^.

| Campaign | *n* | mean pred. | mean actual | signed err. | \|err\| | Spearman |
|---|---|---|---|---|---|---|
| deepseek-r1:8b | 1528 | 0.772 | 0.757 | -0.015 | 0.038 | +0.798<sup>***</sup> |
| llama3.1:8b | 1691 | 0.736 | 0.702 | -0.034 | 0.071 | +0.507<sup>***</sup> |
| llama3.1:70b | 2052 | 0.757 | 0.740 | -0.017 | 0.041 | +0.791<sup>***</sup> |

**Figure 10:**
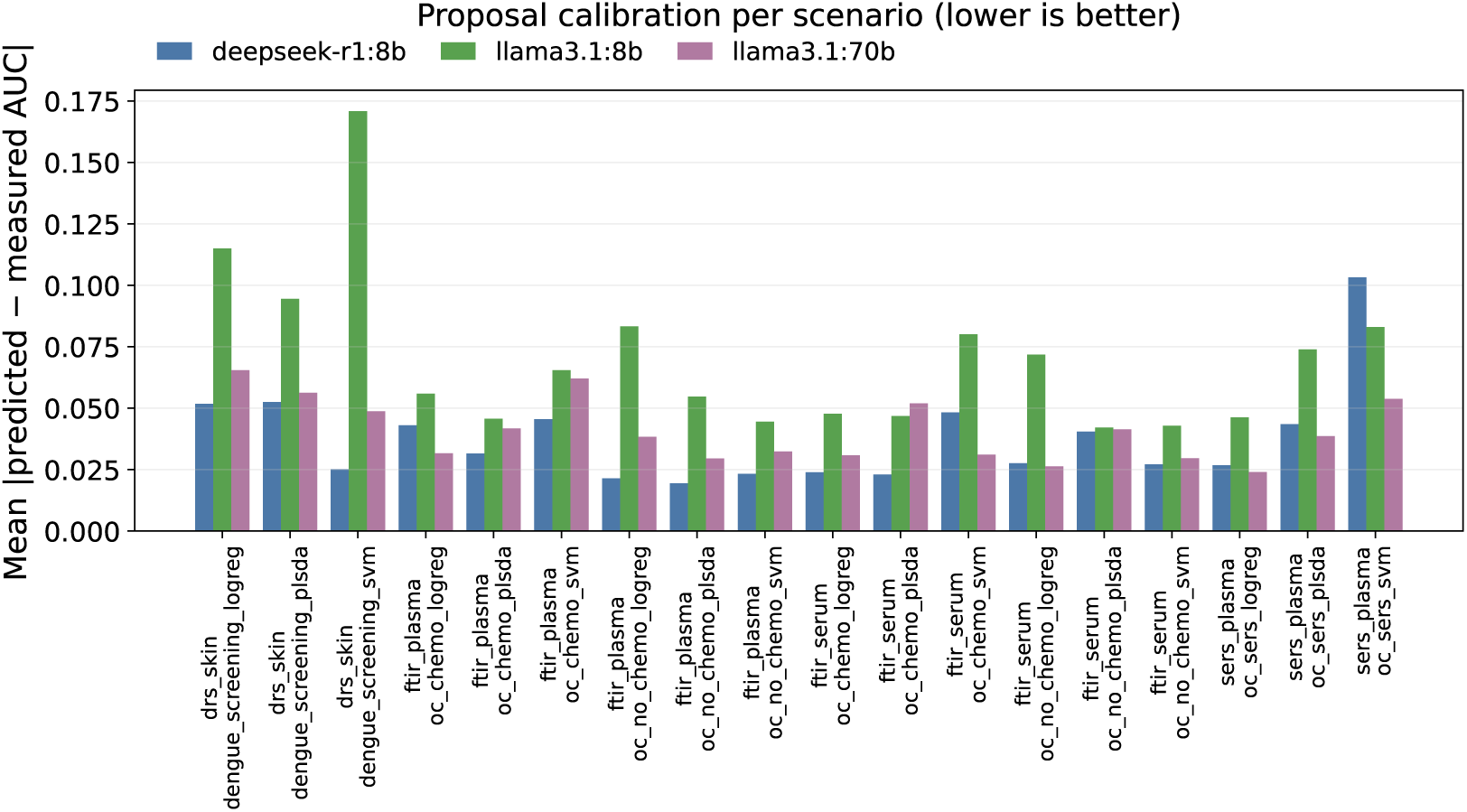
Mean absolute error between the AUC a campaign predicted for a pipeline and the AUC it measured, per scenario. Lower is better. The `llama3.1:70b` campaign improves on `llama3.1:8b` in all 18 scenarios.

**Table 15:** Signed prediction error by method family, pooled over all 18 scenarios. Entries are measured minus predicted AUC, so a negative value is over-prediction, and *±* is the standard error over the *n* proposals of that family. A dash marks a family a campaign never proposed. Averaged over campaigns, a polynomial baseline carries a signed error of +0.002 while every penalised least-squares alternative is over-predicted, at *−*0.019 for `PenalisedPoly`, *−*0.021 for `iModPoly`, *−*0.029 for `IRSQR`, *−*0.044 for `IASLS` and *−*0.050 for `ASLS`.

| Method family | deepseek-r1:8b | llama3.1:8b | llama3.1:70b |
|---|---|---|---|
| WhitakerHayes → SavGol → Poly → Vector | +0.009 ± 0.003 (*n*=62) | −0.019 ± 0.006 (*n*=156) | +0.000 ± 0.002 (*n*=291) |
| WhitakerHayes → SavGol → Poly → SNV | −0.003 ± 0.010 (*n*=6) | −0.018 ± 0.005 (*n*=155) | +0.004 ± 0.002 (*n*=160) |
| WhitakerHayes → SavGol → IASLS → Vector | +0.002 ± 0.012 (*n*=7) | −0.092 ± 0.007 (*n*=228) | −0.061 ± 0.006 (*n*=68) |
| WhitakerHayes → SavGol → IASLS → SNV | −0.058 ± 0.030 (*n*=7) | −0.085 ± 0.009 (*n*=142) | −0.054 ± 0.006 (*n*=106) |
| WhitakerHayes → SavGol → PenalisedPoly → Vector | −0.028 ± 0.014 (*n*=2) | −0.051 ± 0.015 (*n*=63) | −0.020 ± 0.005 (*n*=90) |
| None → Whittaker → Poly → Vector | −0.004 ± 0.004 (*n*=81) | +0.066 ± 0.060 (*n*=2) | −0.010 ± 0.005 (*n*=66) |
| WhitakerHayes → Whittaker → Poly → SNV | −0.007 ± 0.003 (*n*=115) | +0.015 ± 0.013 (*n*=25) | +0.000 ± 0.009 (*n*=8) |
| WhitakerHayes → None → Poly → Vector | +0.017 ± 0.007 (*n*=12) | +0.031 ± 0.016 (*n*=4) | +0.017 ± 0.004 (*n*=113) |
| WhitakerHayes → Whittaker → iModPoly → Vector | −0.027 ± 0.004 (*n*=111) | −0.020 ± 0.014 (*n*=4) | −0.015 (*n*=1) |
| WhitakerHayes → SavGol → PenalisedPoly → SNV | — | −0.068 ± 0.015 (*n*=57) | −0.009 ± 0.005 (*n*=57) |
| WhitakerHayes → SavGol → ASLS → SNV | −0.041 ± 0.037 (*n*=3) | −0.078 ± 0.016 (*n*=46) | −0.030 ± 0.007 (*n*=57) |
| None → Whittaker → IASLS → Vector | −0.032 ± 0.006 (*n*=51) | +0.006 ± 0.033 (*n*=5) | −0.022 ± 0.009 (*n*=47) |
| None → SavGol → Poly → Vector | −0.017 ± 0.024 (*n*=19) | +0.054 ± 0.054 (*n*=3) | −0.015 ± 0.004 (*n*=75) |
| WhitakerHayes → None → Poly → None | −0.019 ± 0.005 (*n*=95) | — | −0.026 ± 0.007 (*n*=2) |
| WhitakerHayes → SavGol → IRSQR → Vector | −0.071 ± 0.031 (*n*=15) | −0.026 ± 0.011 (*n*=65) | +0.010 ± 0.008 (*n*=16) |
| WhitakerHayes → None → PenalisedPoly → Vector | −0.017 ± 0.008 (*n*=6) | +0.069 ± 0.006 (*n*=2) | −0.008 ± 0.006 (*n*=85) |

## E Additional results

**Table 16:** Evaluations by origin, pooled over all 18 scenarios. *Identity* is the all-None pipeline evaluated once per scenario as a reference, and *parametric sampler* is the fallback draw of Algorithm 1 line 11 from the randomly parameterised seed pool.

| Campaign | Origin | *n* | mean AUC | median | max |
|---|---|---|---|---|---|
| deepseek-r1:8b | LLM proposal | 1528 | **0.757** | 0.781 | 0.929 |
|  | Parametric sampler | 614 | 0.648 | 0.658 | 0.904 |
| llama3.1:8b | LLM proposal | 1691 | **0.702** | 0.725 | 0.889 |
|  | Parametric sampler | 451 | 0.665 | 0.672 | 0.887 |
| llama3.1:70b | LLM proposal | 2052 | **0.740** | 0.760 | 0.891 |
|  | Parametric sampler | 90 | 0.688 | 0.704 | 0.878 |
| — | Identity (raw) | 54 | 0.634 | 0.658 | 0.806 |

**Figure 11:**
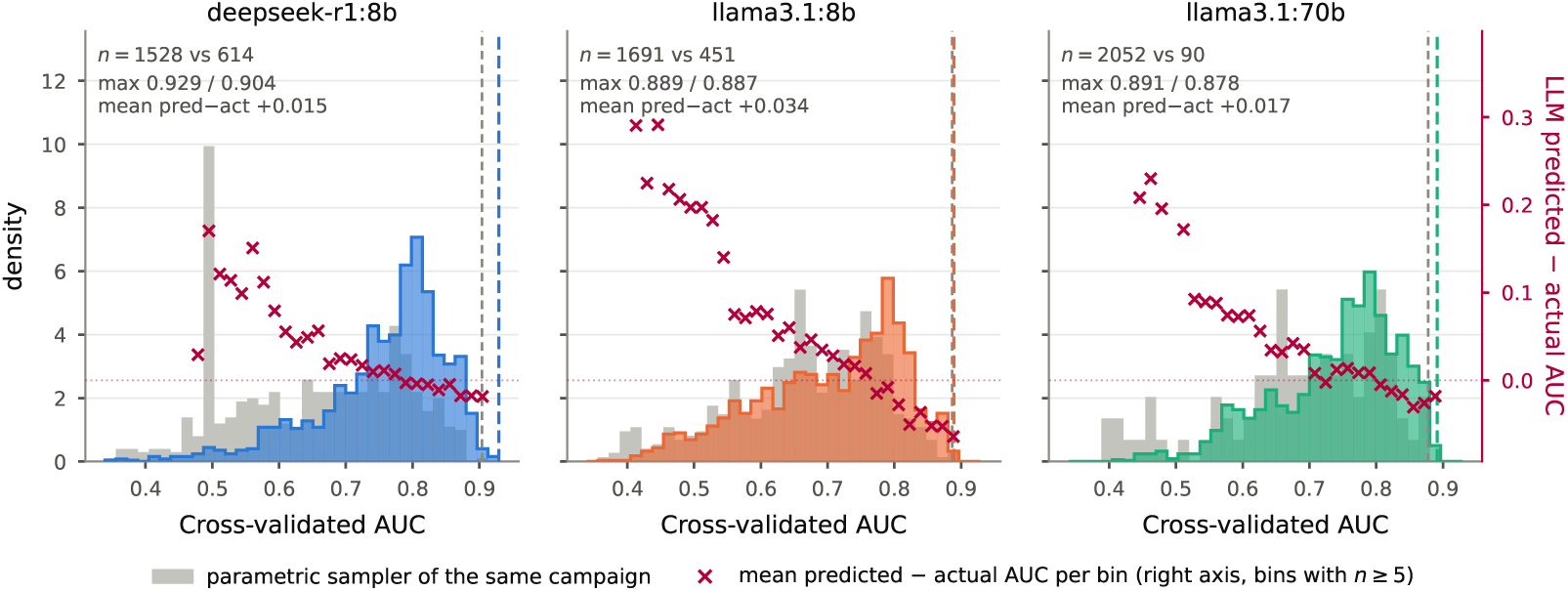
Evaluated pipelines by origin, per campaign (18 scenarios). LLM proposals are shown against the fallback draws of the same campaign (*parametric sampler*), with the mean predicted*−*actual AUC per bin on the right axis.

**Table 17:** Admitted pipelines running a method in a stage it is not registered for. Metrics show totals across two repeats. The AUC comparison is paired within scenario, because misplacements concentrate in particular scenarios and between-scenario AUC differences dwarf the effect. *Scenarios improved* counts the scenarios in which the the averaged AUC of the misplaced pipelines outscored the conformant ones. Most violations arise from methods with similar names (e.g. `Whittaker` denoising placed in the despike stage instead of `WhitakerHayes`) or from closely related penalised least-squares methods used across stages (e.g. `ASLS`/`IASLS` baselines as smoothers since they are closely related to `Whittaker` denoising), as the stage of each method is given in the prompt but not enforced by the validator.

|  | deepseek-r1:8b | llama3.1:8b | llama3.1:70b |
|---|---|---|---|
| admitted pipelines | 4168 | 3716 | 4049 |
| with a foreign family | 66 | 51 | 0 |
| share | 1.6% | 1.4% | 0.0% |
| scenario-splits affected | 18/36 | 17/36 | 0/36 |
| selected pipeline affected (Table 1, 5, and 6) | 1/36 | 0/36 | 0/36 |
| *occurrences, by family and the stage it occupied* |  |  |  |
| Whittaker in baseline (registered: denoise) | 39 | — | — |
| SNV in baseline (registered: normalise) | 14 | 12 | — |
| Whittaker in despike (registered: denoise) | 7 | 15 | — |
| ASLS in denoise (registered: baseline) | 3 | 12 | — |
| IASLS in denoise (registered: baseline) | — | 9 | — |
| Vector in baseline (registered: normalise) | 5 | 2 | — |
| iModPoly in denoise (registered: baseline) | — | 2 | — |
| SavGol in baseline (registered: denoise) | 1 | — | — |
| *effect on CV AUC (training 80%)* |  |  |  |
| paired Δ within scenario-split | −0.026 | −0.005 | — |
| *effect on held-out AUC* |  |  |  |
| paired Δ within scenario-split | −0.047 | −0.011 | — |

**Table 18:** The only miscategorised pipeline that was selected as the best one for a given agent during an agentic run is found on ATR-FTIR serum - no chemo. This is not the best for this scenario overall, see Table 6.

| Stage | deepseek-r1:8b |
|---|---|
| despike | WhitakerHayes<br>kernel_size=7, threshold=15 |
| denoise | **ASLS**<sup>‡</sup><br>diff_order=2, lam=1e+05 |
| baseline | *identity* |
| normalise | SNV |
| CV AUC | 0.828 |
| held-out AUC | 0.819 |
| found in round | 17 |

**Figure 12:**
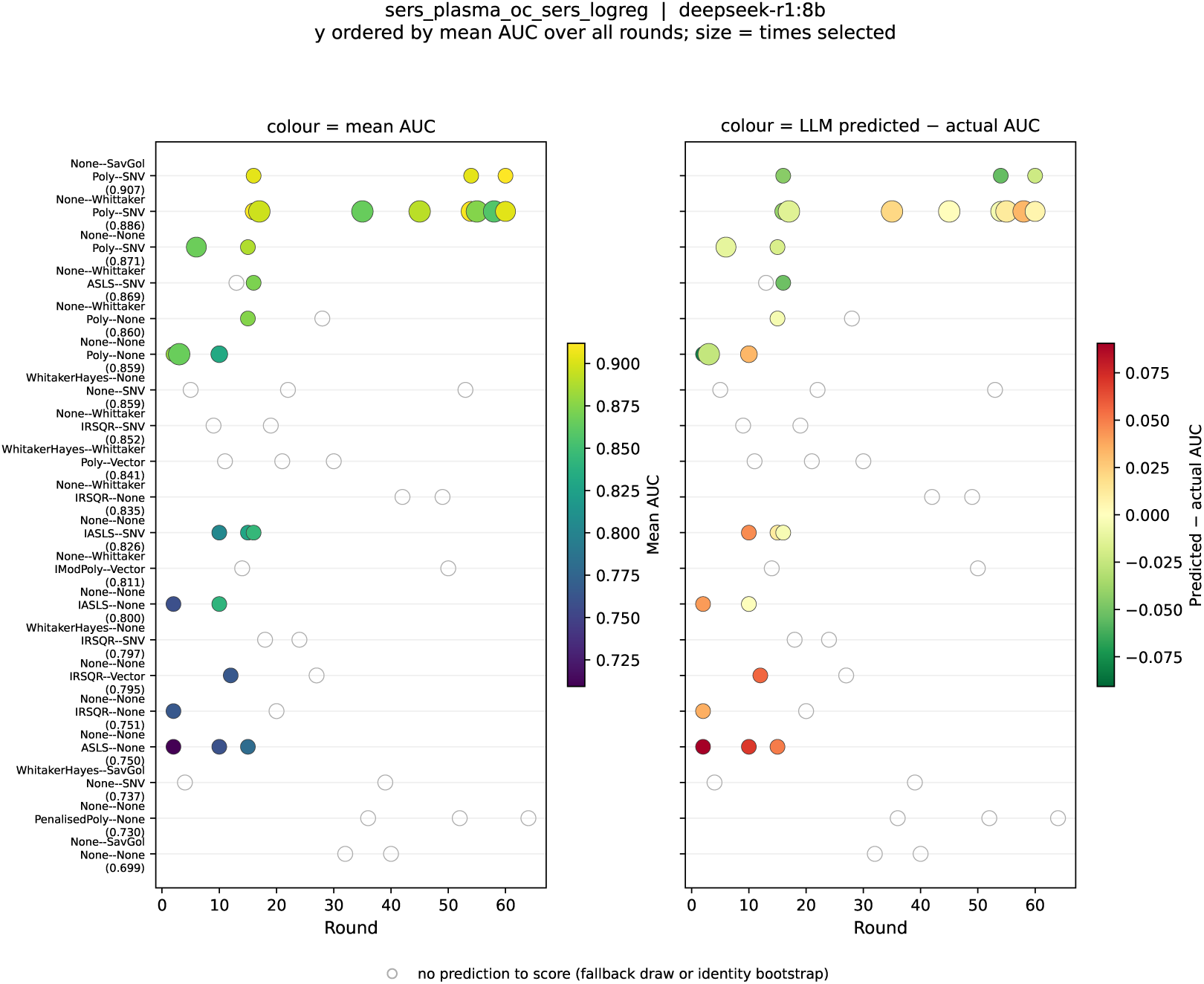
Method families evaluated per round on `sers_plasma_oc_sers_logreg`, `deepseek-r1:8b` campaign (120 evaluations over 64 rounds, 52 distinct families, of which the 20 most-evaluated are shown, covering 73% of evaluations). Round runs against family, bubble area is proportional to the evaluations that family received in that round, and colour and row order are both given by its mean AUC over all rounds. The square at the left margin marks each family’s mean on the same colour scale, so the ordering is legible for families evaluated in a single round. In the RHS figure, colour is the predicted LLM AUC - actual AUC.

**Figure 13:**
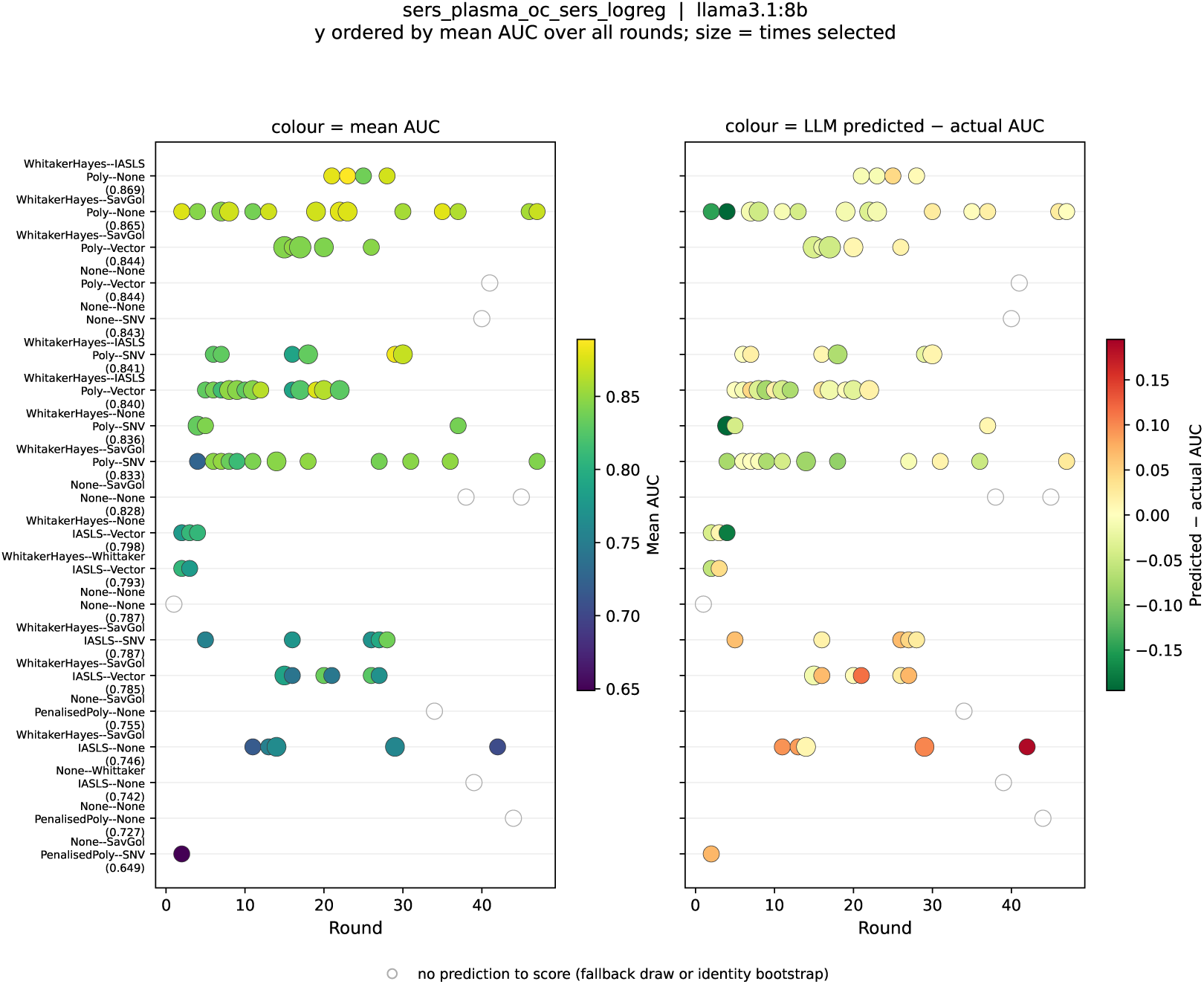
The same scenario under the `llama3.1:8b` campaign (120 evaluations over 49 rounds, 30 families, of which the 20 most-evaluated are shown, covering 92% of evaluations).

**Figure 14:**
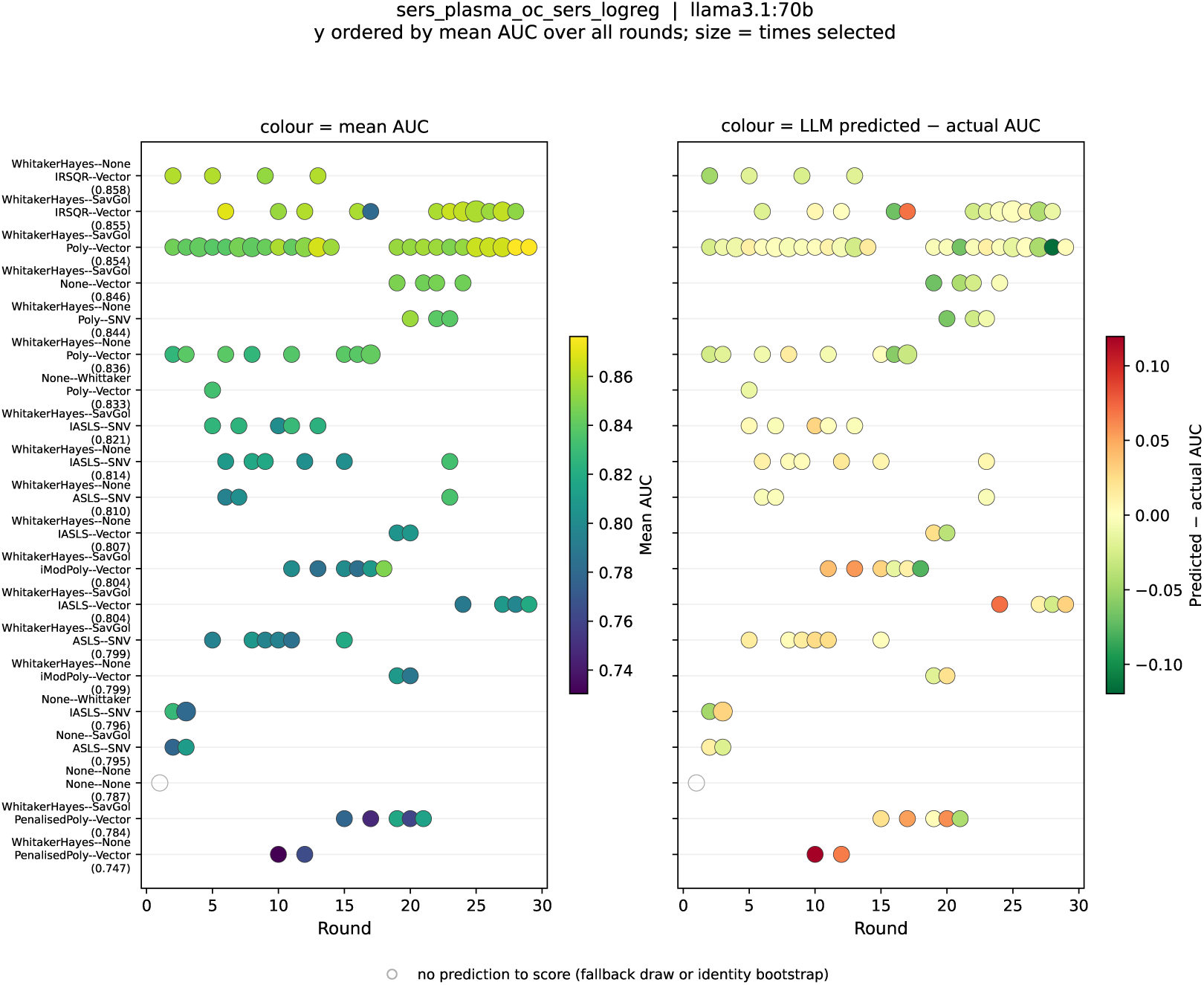
The same scenario under the `llama3.1:70b` campaign (120 evaluations over 29 rounds, 24 families, covering 97% of evaluations).

**Table 19:**
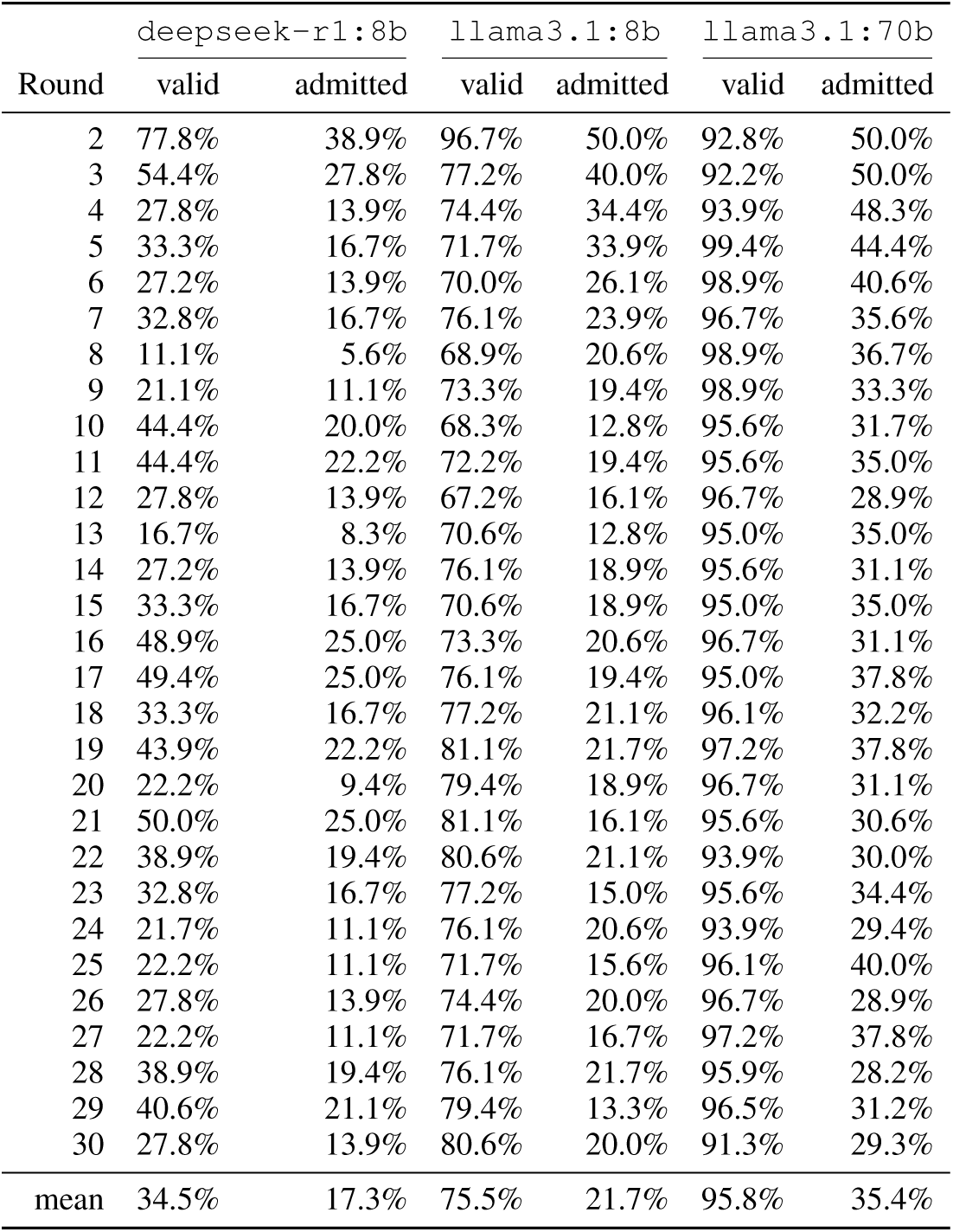
Proposal yield per round, pooled over all 18 scenarios. The denominator is what the search asked for, *K* = 10 proposals per scenario-round, not what came back parseable, so a round whose response could not be extracted contributes zeros rather than being dropped. *Valid* is the share passing the §5 bounds and §8 schema checks. *Admitted* the share surviving those checks and de-duplication against the existing action space. Admission is capped at *C*_grow_ = 5 new combinations per round, so no cell can exceed 50%. The early rounds of both `llama3.1` campaigns sit exactly at that ceiling and are censored, not merely high. Only scenario-rounds in which a request was issued are counted, so a campaign that exhausts its budget early is not charged for rounds it never ran.

